# TGFβ−induced embryonic cell senescence at the origin of the Cornelia de Lange syndrome

**DOI:** 10.1101/2022.07.26.501526

**Authors:** Céline Hachoud, Faten Chaabani, Erwan Watrin, Manuela Wuelling, Heiko Peters, Valérie Cormier-Daire, Michel Pucéat

**Affiliations:** INSERM U1251 Aix-Marseille University, Marseille, France; Centre National de la Recherche Scientifique, UMR6290, Rennes, France; Institut de Génétique et Développement de Rennes, Université de Rennes, Rennes, France; Centre for Medical Biotechnology, Developmental Biology, University Duisburg- Essen; Newcastle University Biosciences Institute, Centre for Life, Newcastle, NE1 3BZ, UK; Université Paris Cité, INSERM UMR 1163, Institut Imagine, Paris, France; Service de médecine génomique des maladies rares, Centre de Référence Pour Les Maladies Osseuses Constitutionnelles, AP-HP, Hôpital Necker-Enfants Malades, Paris, France; CoHEART Consortium collaborators: Gregor Andelfinger, Jeroen Bakkers, Bart Loeys

## Abstract

Cornelia de Lange Syndrome (CdLS) largely caused by mutation of the cohesin loader NIPBL is a rare developmental disorder affecting the formation of many organs. Besides a short body size and neurological defects, more than half of CdLS children feature various cardiac malformations.

To mimic the disease and test a therapeutic strategy, we generated a C57/Bl6 *Nipbl+/-* mouse model of the disease. These mice featured a severe delay in both embryonic and postnatal growth. The *Nipbl-*deficient embryonic and neonatal hearts developed ventricular hypertrophy, aortic and valve defects associated with a persistent truncus arteriosus and a ventricular septal defect. Muscles derived from the second heart field were deficient in the *Nipbl* haplo-insufficient mouse embryos. The adult hearts then featured a severe aortic senescence phenotype and a stenosis resulting in an increase in aortic flux velocity and persistent left ventricular hypertrophy. Using proteomics and RNA-sequencing in embryos, we identified a dysregulated TGFβ pathway in the outflow tract of embryonic hearts as well as the presence of senescent cells as early as in E13.5 *Nipbl+/-* embryonic hearts, limb primordium cartilage as well as in different post-natal tissues including muscle and brain cortex. Treatment of pregnant *Nipbl+/-* mice with a TGFβR (ALK5) inhibitor from E9.5 to E13.5 prevented cell -senescence and rescued the cardiac phenotype as well as the body size of mice at birth.

Altogether our data revealed that an exacerbated TGFβ pathway associated with cell senescence is at the origin of many defects in a CdL mouse model. This druggable pathway opens the path toward a potential preventive and/or therapeutic strategy for post-natal CdLS patients.

## Introduction

Cornelia de Lange syndrome (CdLS) is a rare genetic and developmental disorder affecting about 1:10,000/1:30,000 children. The syndrome is sometimes diagnosed at prenatal stages and most often at birth because of distinct facial features of babies due to cranio-facial malformations. A majority of CdLS children also presents with a short stature, mild to profound neuro-cognitive disabilities, microcephaly, upper limb defects, as well as gastro-esophageal reflux^1,2^.

Cardiac defects are also observed in more than 50 % of patients^3,4^. Cardiac malformations arise in principle from defects in differentiation, and/or migration of cardiac progenitors and of cells emerging from the embryonic second heart field^5^. CdLS children hearts feature septal defects and outflow tract defects including hypoplastic aorta, stenosis, or coartation of great arteries as well as Tetralogy of Fallot.

Since its discovery, CdLS has been described as a clinically highly variable disease, which suggested a multigenic origin or a causative gene playing a multifunctional role.

The first and main *NIPBL* (Nipped-B-like protein) gene whose mutations are responsible for the syndrome has been uncovered in 2004^6^. NIPBL main function is to load the cohesin complex onto DNA and to ensure a stability of the genome^7^ although NIPBL may also work like a transcription factor binding gene regulatory regions independently from cohesin^8^.

Although other genes encoding proteins of the cohesin complex or associated proteins (*RAD21, SMC1A, SMC3, HDAC8, BRD4)* have also been found mutated in CdLS patients or in patients with associated syndromes, *NIPBL* mutations and in turn gene haploinsufficiency explain a large spectrum of CdLS patients^9^. While the syndrome has been described in 1933 by Pr Cornelia de Lange^10^, no therapeutic strategy has been proposed and patients have to be managed by clinicians on a symptomatic and social basis. Both the clinical heterogeneity of patients and the likely multiple functions of a large multi-domain protein such as NIPBL make both the understanding of the disease and the research of therapeutic targets challenging.

Several cell^11–13^ and animal models including zebrafish and mice^13–18^ have been developed to better understand the disease and to reach such a therapeutic aim. However, most of these models do not faithfully recapitulate the human syndrome at least in a reproducible manner. They however allowed to identify dysregulation of specific genes and of pleiotropic signaling pathways such as the Wnt pathway^11,13^ as well as DNA repair and cell senescence pathways^19–21^ in *nipbl* haploinsufficient cells but no therapeutic strategy could emerge from these studies.

Interestingly, a crosstalk was recently described between cohesinopathies and TGFβ-related disorders^22^. For example, CdLS patients feature some defects in great arteries that are common to those observed in Marfan or Loeys-Dietz syndrome^23^. More specifically, TGFβ-dependent pathway was found to be constitutively activated in a recently described cohesinopathy^24^. Interestingly, SMC protein (SMC3), a cohesin complex component also named chondroitin sulfate proteoglycan 6 is an extracellular protein expressed in smooth muscle cells and activated by TGFβ^25^. Interestingly, SMC3 mutated CdLS patients feature a high incidence of cardiac defects^26^, Being aware of mouse models of genetic diseases that may or not recapitulate human diseases according to their genetic background and overall level of expression of the gene of interest ^27,28^, we first characterized the cardiac phenotype of a novel C57Bl/6J mouse model of *Nipbl* haplo-insufficiency and combined it with human CdLS patient-specific iPS cells in order to uncover a potential therapeutic target of CdLS.

We found that *Nipbl+/-* mice featured a significant decrease in *Nipbl* mRNAs as well as a moderate decrease in the protein in the heart. These mice also present a severe delay in embryonic and postnatal growth. The heart at birth featured ventricular hypertrophy associated with a persistent truncus arteriosus (PTA). The adult hearts then feature a severe aortic phenotype with an enlargement of the intima including senescent cells and a stenosis resulting in an increase in aortic flux velocity and persistent left ventricular hypertrophy. Using proteomics and RNA-sequencing, we identified a dysregulated TGFβ pathway in the outflow tract of embryonic hearts and the presence of senescent cells as early as in E13.5 *Nipbl+/-* embryonic hearts, as well as in post-natal muscle and neonatal brain cortex.

Treatment of pregnant mice with a TGFβR (ALK5) inhibitor from E9.5 to E13.5 prevented cell senescence and rescued the cardiac phenotype as well as the body size of mice at birth.

## Results

### Characterization of Nipbl+/- mice

*Nipbl+/-* haplo-insufficient mice were generated by deleting the exon 2 that includes the ATG of *Nipbl* in one allele using an ubiquitous CAG*^cre^* mouse and the *Nipbl floxed* mouse^29^. The *Nipbl+/-* mice were then backcrossed for ten generations in the C57Bl/6J genetic background.

C57Bl/6J *Nipbl+/-* mice featured a severe growth delay as shown by the small size of E13.5 embryos, neonates as well as 2 months old adult mice (Fig. 1a). In order to better evaluate this growth delay, we first measured the length of the tibia, the thickness of ribs as well as the length of fingers in the front leg of neonatal mice in skeleton stained with Alizarin Red and Alcian Blue. Figure 1b and 1c revealed a significant decrease in all measured parameters in *Nipbl+/-* mice compared to wild type (wt) mice. We then more specifically investigated the radius composition in neonatal mice at the cellular level. Both the radius and its hypertrophic zone were significantly shorter in *Nipbl+/-* mice when compared to *wild type* (Fig.1d). The reduced hypertrophic zone was associated with a reduced expression domain of Collagen type 2 *(Col2)* and *10a1 (Col10a1)* in *Nipbl+/-* mice when compared to *wild type* littermates (Fig. S1).

**Fig. 1.**
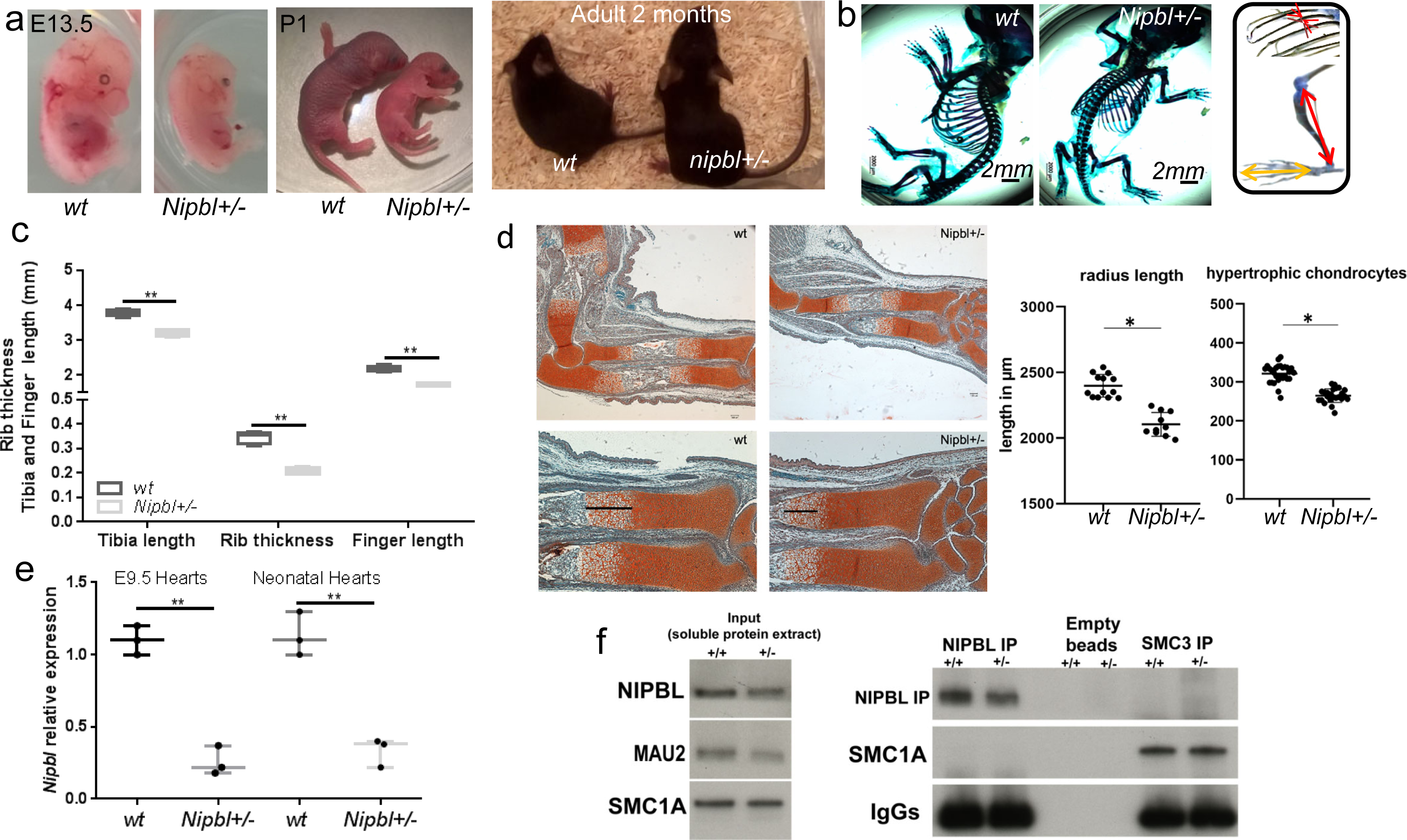
Characterisation of *Nipbl+/-* mouse. **a,** from left to right: E13.5 embryos, neonates and 2 months adult mice. The images are representative of at least 15 embryos or mice at each stage of development. **b,** Alizarin red and Alcian Blue stained neonatal skeletons. **c,** Graph of the size of tibia, the thickness of ribs and the length of fingers (measurements made as shown in the right inset) (n=5, ** p<0.01). **d,** Total radius length and the hypertrophic zone (black line) were measured in Safranin-Weigert stained forelimbs sections of E16.5 Nipbl+/- and wt mice. Dots show individual measurements of serial sections from 2 Nipbl+/-, or 2 wt embryos,* p<0.05, scale bar: 100 µm). **e,** Q-PCR of *nipbl* from E9.5 embryonic hearts and neonatal hearts (n=3 from 3 separate litters;** p<0.01), **f,** Western blot of NIPBL using an antibody directed against the N-terminal domain of NIPBL (Bethyl) and SMC1A in whole lysate of neonatal heart or after immunoprecipitation using anti-whole NIPBL antibody (Abcam)

We next monitored expression of *Nipbl* mRNA in E9.5 and neonatal whole *Nipbl+/-* and wild type hearts by Q-PCR. Figure 1e shows a significant decrease (down to 20 %) in *Nipbl* transcripts in *Nipbl+/-* when compared to *wild type*. We also looked at the protein level more specifically in cardiomyocytes isolated and purified from neonatal hearts. Western blot (direct or after immunoprecipitation) also showed a decrease of about 30 % confirmed by blotting after enrichment by immunoprecipitation of the NIPBL protein. Interestingly, Mau2 was also decreased in *Nipbl+/-* haploinsufficient myocytes (Fig.1f) while cohesin complex subunit SMC1A was unaffected.

### Cardiac phenotype of *nipbl+/-* mice

We first looked at the cardiac phenotype of *Nipbl+/-* mice at birth. High Resolution Episcopic Microscopy (HREM) revealed that *Nipbl+/-* mice featured a severe ventricular hypertrophy (Fig.2a). The wall of left ventricle was twice as thick in *Nipbl+/-* mice when compared to *wt* (Fig.2b). This hypertrophy was associated with a septation defect of the great vessels and in turn, both a persistent truncus arteriosus and a stenosis of the distal artery in *Nipbl+/-* hearts. The aortic valve leaflets were also thickened in *Nipbl*+/- hearts when compared to wild type hearts (Fig 2a) as further revealed by Mowat staining (Fig.2c). The right ventricle, which usually appears as crescent shaped in *wt* hearts, largely lost this shape in *Nipbl+/-* hearts (Fig.S2). The latter also revealed a ventricular septum defect (i.e., an interventricular communication) at the apical region (Fig.S2).

**Fig. 2.**
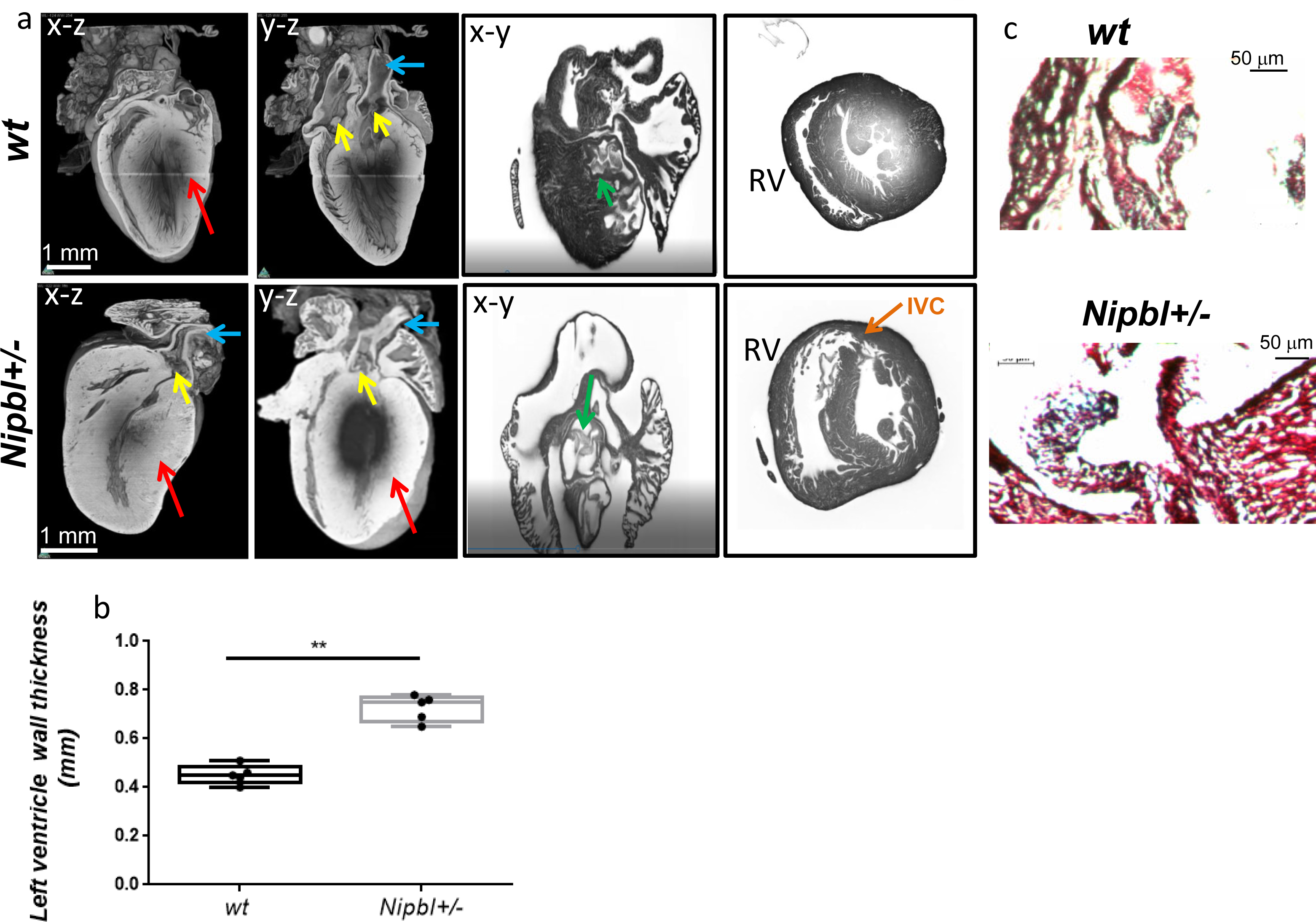
Phenotype of neonatal *Nipb*l+/- mouse hearts. a, HREM images of x-z, y-z, and x-y sections of wt and *Nipbl+/-* neonatal hearts . The red arrows point to the thickness of the left ventricular wall; the yellow arrows point to the PTA in *Nipbl+/-* heart; the green arrows point to the aortic valve. The blue arrows point to a stenosis of distal aorta in *Nipbl+/-* heart (left panels) and the orange arrow to an interventricular communication (IVC) in right pannels **b,** Thickness of the compact wall of the left ventricles of wt and n*ipbl+/-* neonatal hearts (n=5); t-test **p<0.01. **c,** Mowat staining of aortic valves.

Cardiac hypertrophy was also observed in two months old adult *Nipbl+/-* mice first revealed by an increase in contractility of left ventricle as monitored in echocardiography (Fig.3a). Doppler Echocardiography also showed an increase in aortic flux velocity (Fig.3b) associated with a decrease in the diameter of the aorta (Fig.3c) of *Nipbl+/-* mice when compared with *wild type* mice. The phenotype was fully penetrant and observed in 92% of mice (n=24).

Immunostaining of hearts and more specifically of the aorta with an anti-smooth muscle actin (SMA) antibody showed an increase in thickness of the aortic wall of *Nipbl+/-* mice (Fig.3d-e) when compared to wild type mice. Interestingly the media of *Nipbl+/-* mice aorta was filled with large cells which did not express SMA (Fig.3d yellow inset). Of important note, these cells were positive for γH2AX suggesting a senescent phenotype.

Conditional deletion of *Nipbl* specifically in the smooth muscle lineage using the SMA*^CreERT^*^2^ mouse with the recombinase activated by tamoxifen at E11.5 fully recapitulated the functional aortic phenotype observed in *Nipbl+/-* mice. As a consequence, adult mice lacking *Nipbl* in smooth muscle cells featured an increased aortic flux at 4, 7 and 11 weeks (Fig. S3).

### Skeletal muscle phenotype of *nipbl+/-* mice

We next looked at the skeletal muscles derived from the second heart field^30,31,32^ in both *Nipbl+/-* mice and after specifically deleting *Nipbl* in the second heart field using the *Mef2cAHF^cre^* mouse. *In Mef2cAHF^cre^/Nipbl^fl/fl^*homozygous E13.5 embryos, the trapezius muscles, the extra-ocular muscles, the oesophagal muscle, the pervertebral muscles of the neck and the masseters were atrophied or missing when compared to heterozygous *Mef2cAHF^cre^/Nipbl^fl/+^* embryos (Fig.S4). Similar observations were recorded in *Nipbl+/-* embryonic mice (data not shown).

As many CdLS patients present gastro-oesophagal reflux, we further examined the skeletal oesophagal muscle in 2 months old adult mice and found that this muscle was markedly less developed in *Nipbl+/-* mice (thickness 19±3 μm, n=3) than in *wild type* mice (47±5 μm, n=3) (Fig. S5a).

### Senescence at the origin of tissue and organ defects in *Nipbl*+/- mouse

In order to characterize the origin of PTA we investigated back the embryonic heart. *Nipbl*+/- haplo-insufficient embryonic E13.5 hearts (n=9 from 4 separate litters) also featured the septation defect of the outflow tract (Fig. 4). Interestingly we found that many p21+ senescent cells accumulated at the base of the common outflow vessel. We found 1.68±0.11 % (n=5) of p21+ cells in wt hearts versus 104±16 % (n=5) in *Nipbl+/-* hearts. Furthermore, five times more cells were positive for P-ERK in the same area in *nipbl+/-* heart than in wt hearts (Fig. 4, Fig.S5). Forelimb primordium cartilage of wt E13.5 embryos featured spatially located areas of p21 + cells. In contrast, *Nipbl+/-* embryonic primordium cartilage revealed a very large area of p21+ cells (Fig. 4c). Similarly, we looked back to the oesophagus skeletal muscle. We found that each 10 μm section of *wt* muscle cell featured one or none p21+ cells while the ones of *Nipbl+/-* mice revealed 8±2 p21+ cells (n=3) (Fig.S5b). We also observed in the cortex of neonatal mouse brains an abnormal presence of p21+ neuronal cells in *Nipbl*+/- mice compared to *wt* (Fig S5d). Altogether, these observations strongly point to senescence as a major consequence of nipbl haploinsufficiency.

**Fig. 3.**
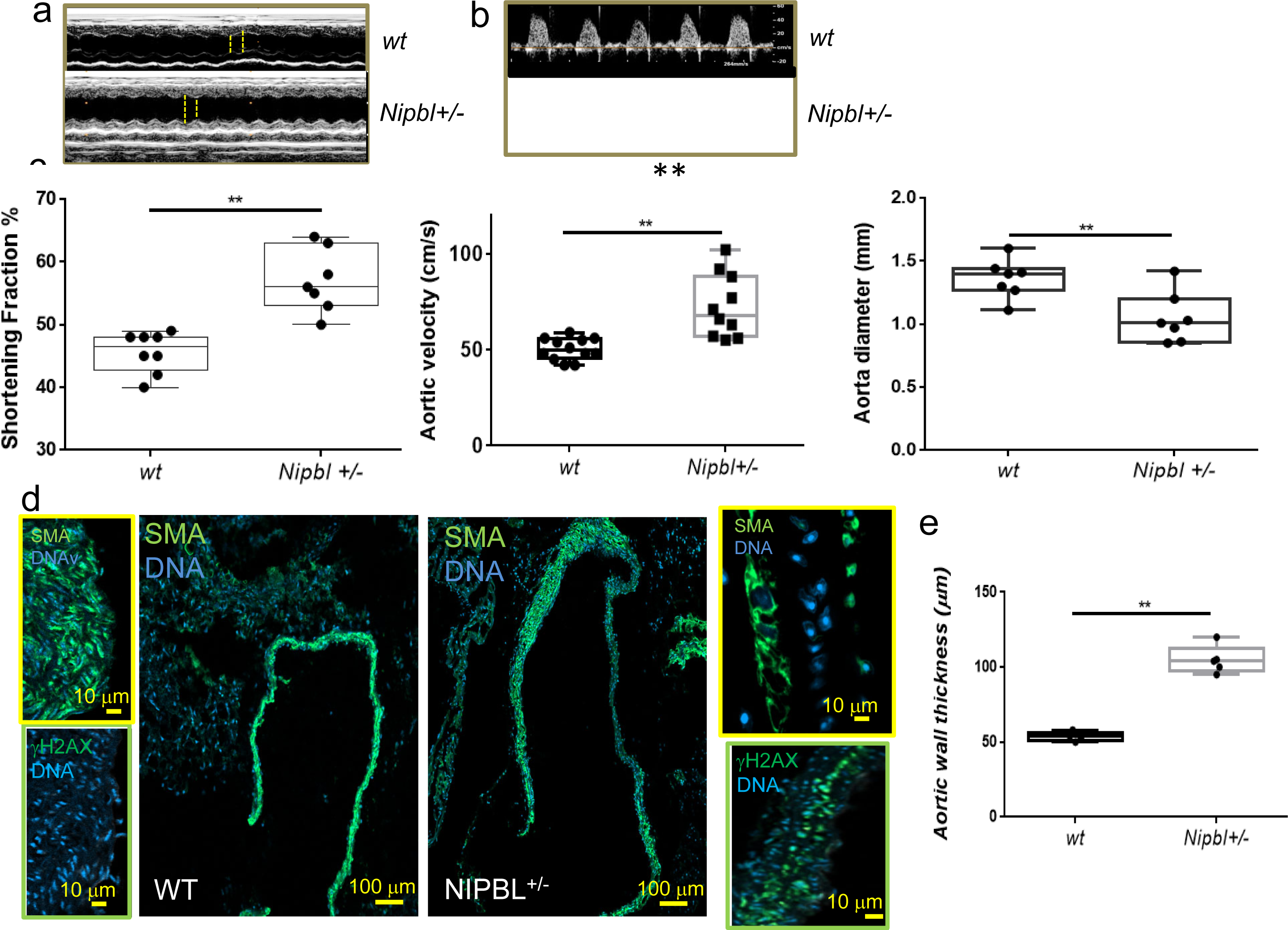
Adult cardiac phenotype of nipbl+/- mice. **a,b,c** echocardiography of 2 months old mice. **a,** shortening fraction **b,** maximal aortic flux velocity **c,** diameter of the aorta (n≥ 7 mice) **d,** Anti–SMA staining of aorta. Yellow insets show high magnification of cells within the aortas. Green inset:the aorta of WT and NIPBl+/- aorta were stained by an anti γH2AX antibody. **e,** graph of aortic wall thickness (n=5) t-test **p<0.001.

**Fig. 4.**
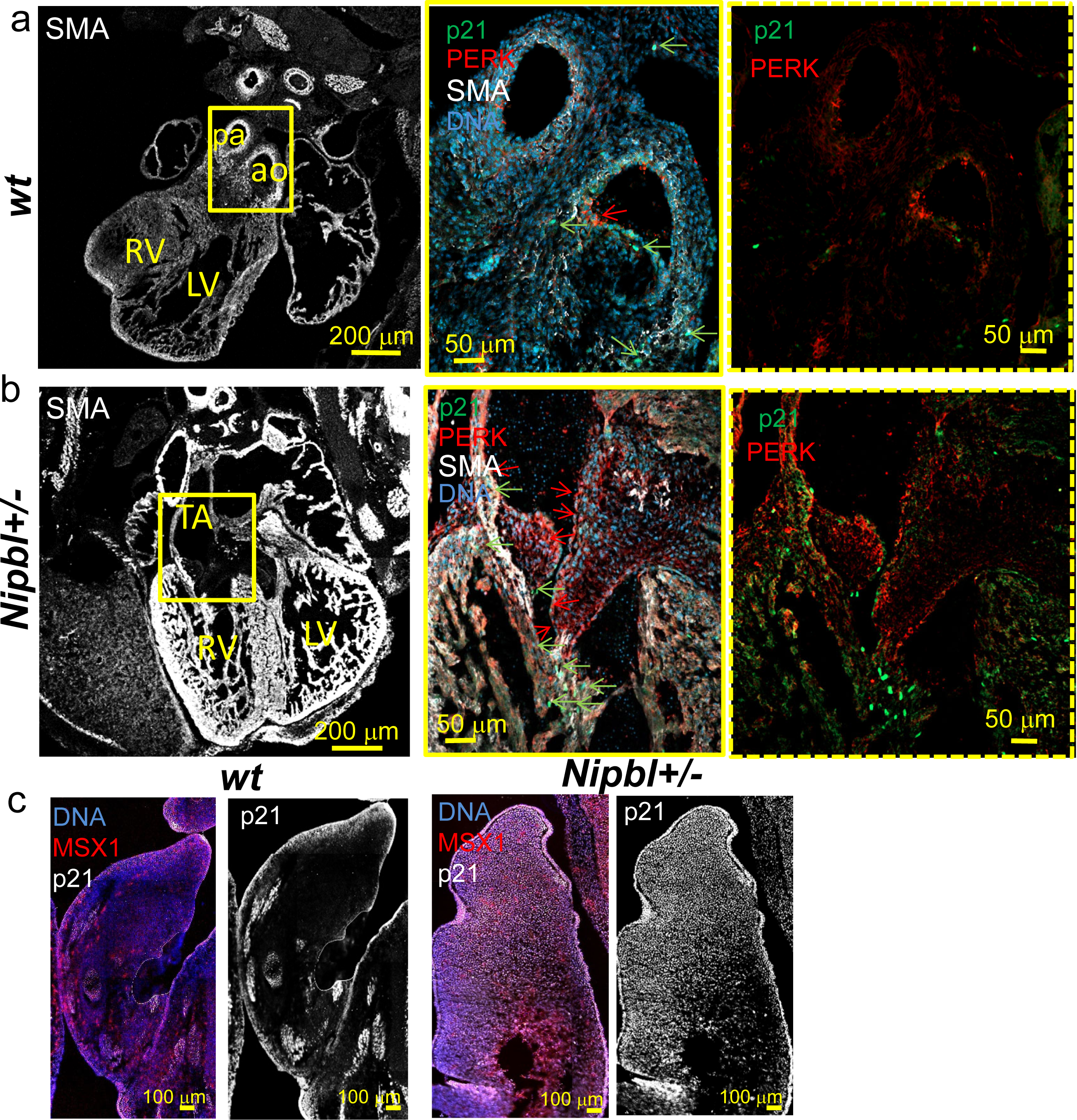
Embryonic sections from *wt* and *Nipbl+/-* E13.5 embryos. Heart sections from wt **(a)** and Nipbl+/- **(b)** embryos were stained with anti-SMA, anti-p21 and anti– pERK antibodies. In wt heart, both aorta and pulmonary trunk (pa) are present. In Nipbl+/- heart, a common outflow vessel is formed. Insets on the right show high magnification of outflows; green arrows point to p21+ nuclei; red arrows point to pERK+ nuclei. The right panel (dashed line) shows only p21 and pERK + nuclei. ao:aorta,pa pulmonary trunk,LV:left ventricle, RV right ventricle,TA truncus arteriosus. (**c**) forelimbs were stained with anti-msx1 and anti-p21 antibodies. (only p21 is shown on right images. The figure is representative of at least 9 embryos from different litters

We thus searched for the origin of cell senescence in *Nipbl*+/- mice. More specifically, to better understand the outflow tract phenotype of *Nipbl+/-* mice, we performed a proteomic analysis of the outflow tract in E16.5 embryos. 6 *wt* and 6 *Nipbl+/-* embryos were collected, the heart explanted and the outflow tract entirely dissected out. Each OFT was individually subjected to mass spectrometry analysis. As shown in Figure 5, *Nipbl+/-* OFT featured a loss of proteins present in the extracellular matrix (*Supplementary data 1*). We specifically highlighted four proteins emelin1, fibrillin 1 and 2 and actbl2 that are dramatically under-expressed in *Nipbl+/-* OFT as compared to that of *wild-type*. Remarkably, all these proteins are involved in the TGFβ signaling pathway including fibrillins, which sequester TGFβ^33^ or in related aortic diseases^23^.

**Fig. 5:**
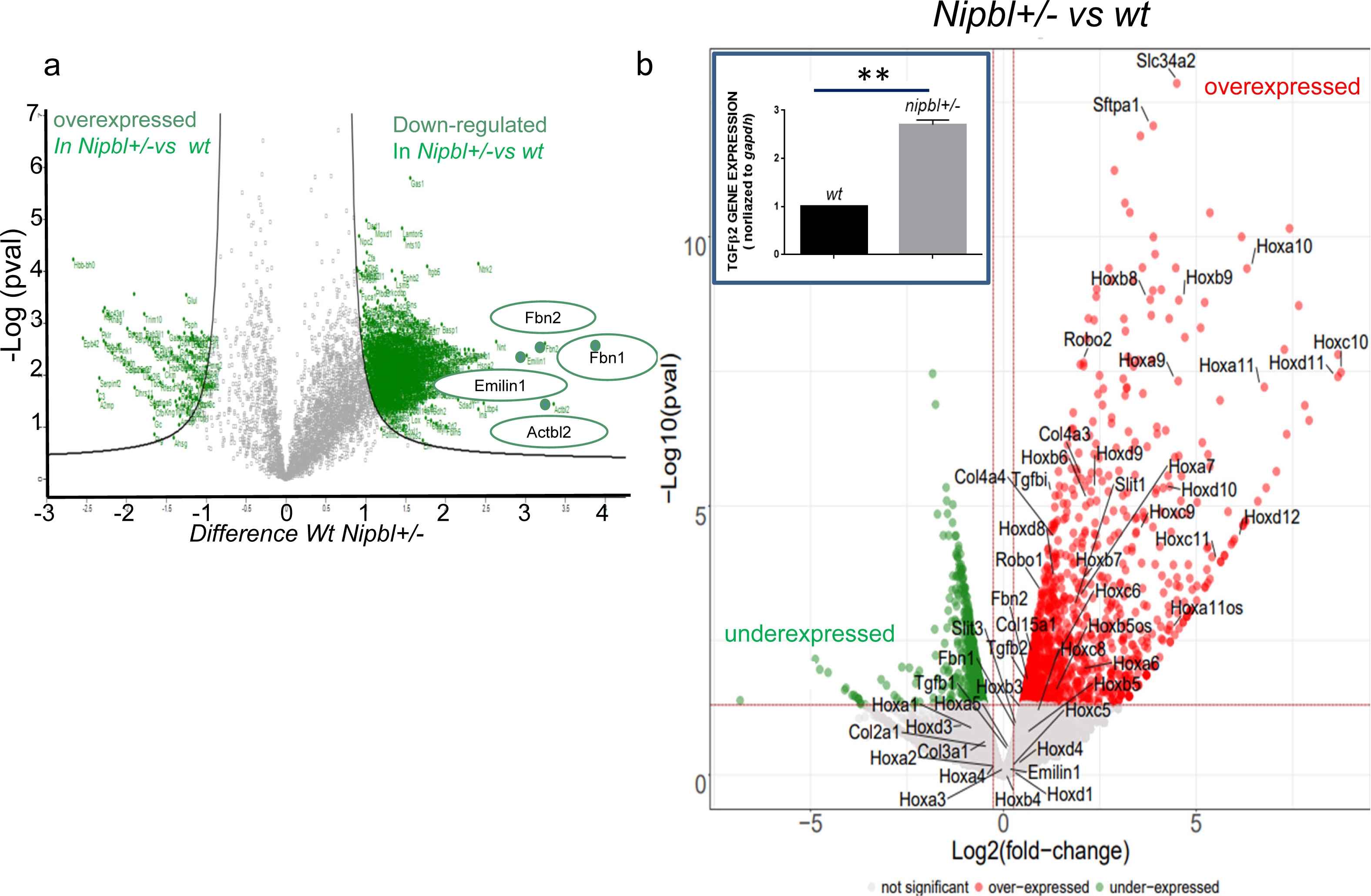
**Proteins and genes expression of OFT of nipbl+/- embryonic hearts**. 12 *wt* and 12 *Nipbl+/-* E16.5 embryos were collected from 6 separate litters, the hearts explanted and the outflow tract entirely dissected out to be processed as separated samples in proteomic analysis (**a**). Volcano plot is shown in **a.** 6 wt and Nipbl+/- OFT were used as bulks from 6 different litters for RNA-sequencing. **b**, Volcano plot of OFT genes dysregulated in *Nipbl+/-* mice Versus wild type. Inset in b: RT-QPCR of TGFβ2 ** n=6 t-test p<0.001

RNA-sequencing of the same cardiac regions (7 OFT from E16.5 *wild type* and *Nipbl+/-* embryonic hearts) revealed an increase in expression of 1199 genes (*Supplementary data 2)* including genes involved in OFT formation such as *slit2, Robo1, six2* and *Hoxb, Hoxc* and *Hoxd* as well as *TGFβ2* genes in *Nipbl+/-* compared to *wt* OFT. TGFβ2 upregulation was further confirmed by RT-Q-PCR (Fig.5b, inset). Anterior *Hoxa* genes were for most of them downregulated in OFT of *Nipbl+/-* vs *wild type* mice as well as *pbx1,* all genes whose haploinsufficiency are associated with PTA^34,35^.

### CdLS iPS cells and smooth muscle derivative feature a cell senescence phenotype

To get more insight into the molecular mechanisms underlying senescence in *Nipbl+/-* haplo-insufficient mice, we switched to a human model using iPS cells derived from CdLS patient cells. 3 iPS cell lines were used from patients harboring the *NIPBL* mutations c.6242.g>C in exon 35, or c.6860T>C in exon 40 or c.6516- 6517 in exon38. All patients featured cardiac malformations. Both the human embryonic stem cell line H9 and an iPS cell line from a healthy volunteer were used as controls. CdLS IPS cells showed a decrease in NIPBL and an exclusion of the protein from the nucleolus. We also observed fragmentations of nucleoli in all CdLS IPS cells (Fig.6 a,b) in contrast to wt cells which showed only two nucleoli, a feature of pluripotent stem cells. Cells were then differentiated in smooth muscle cells. While wild type cells still proliferated after one week of differentiation, CdLS cells stopped dividing and expressed for most of them p21 (Fig.6c). Thus, the analysis of human cells confirmed that CdLS cells are programmed early towards a cell senescence phenotype.

**Fig. 6.**
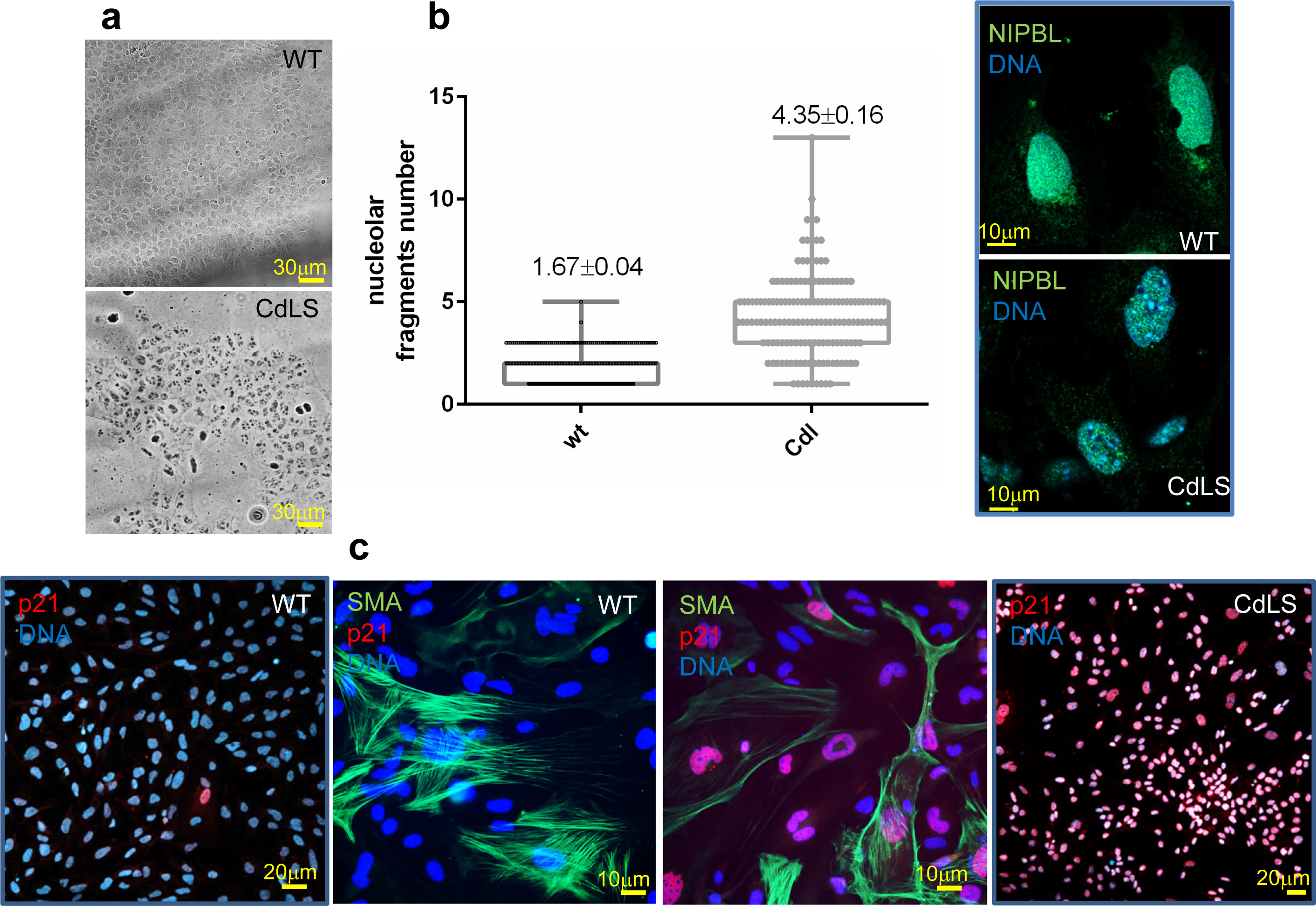
iPS cells from wt or CdlS patient (CdL) **a,** undifferentiated cells (inset on the right: anti-NIPBL stained cells). **b,** graph showing the number of nucleolar fragments /cell. **c,** cells were diferentiated in smooth muscle cells and stained after one week with anti-p21 only (insets) or with both anti-p21 and anti-SMA antibodies . The figure is reprensentative of 3 experiments performed using each of the 3 CdLS iPS cell lines as well as both H9 HUES cell line and a control (wt) iPS cell line

### Decreased motility of mesenchymal cells of proximal OFT

As senescence is known to affect cell motility, and in order to confirm senescence phenotype, we investigated the motility of cells undergoing epithelial-to-mesenchymal transition (EMT) in the proximal outflow tract at E10.5 in both wt and *nipbl+/-* embryonic hearts. To such an aim we cultured explants dissected out from proximal OFTs on collagen gel for 48 hrs. We then monitored the distance of migration of cells undergoing EMT from the explant. Wild type cells migrated significantly farther than *Nipbl+/-* cells (Fig.S6) consistent with a senescence state conferred by *Nipbl* haplo-insufficiency.

### A TGFβ (ALK5) receptor inhibitor rescues the phenotype of nipbl+/- mice

As our data pointed to a hyperactive TGFβ pathway, we reasoned that its inhibition could prevent the senescence and the associated phenotypes observed in *Nipbl* haplo-insufficiency mouse model. Thus, we tested the effect of the ALK5 inhibitor Galunisertib on *Nipbl+/-* mice. Pregnant mice were treated from E9.5 up to E13.5 with 30 mg/kg/day of galunisertib. Embryos were first collected at E13.5. Remarkably, and as illustrated in Figure 7, embryonic *Nipbl+/-* hearts did not feature anymore PTA when compared to wild type hearts from the same litter. Distinct pulmonary trunk originating from the right ventricle and aorta from the left ventricle were indeed observed (Fig. 7a). in addition, very few p21+ cells were observed at the base of the outflow region of both the wild type and *Nipbl+/-* embryonic hearts (inset in Fig. 7a). Expression of both *Col2* and *Col10* was similarly fully rescued (Fig.S8). Such a rescue was already observed at E18.5 stage of embryonic development (data not shown).

**Fig. 7.**
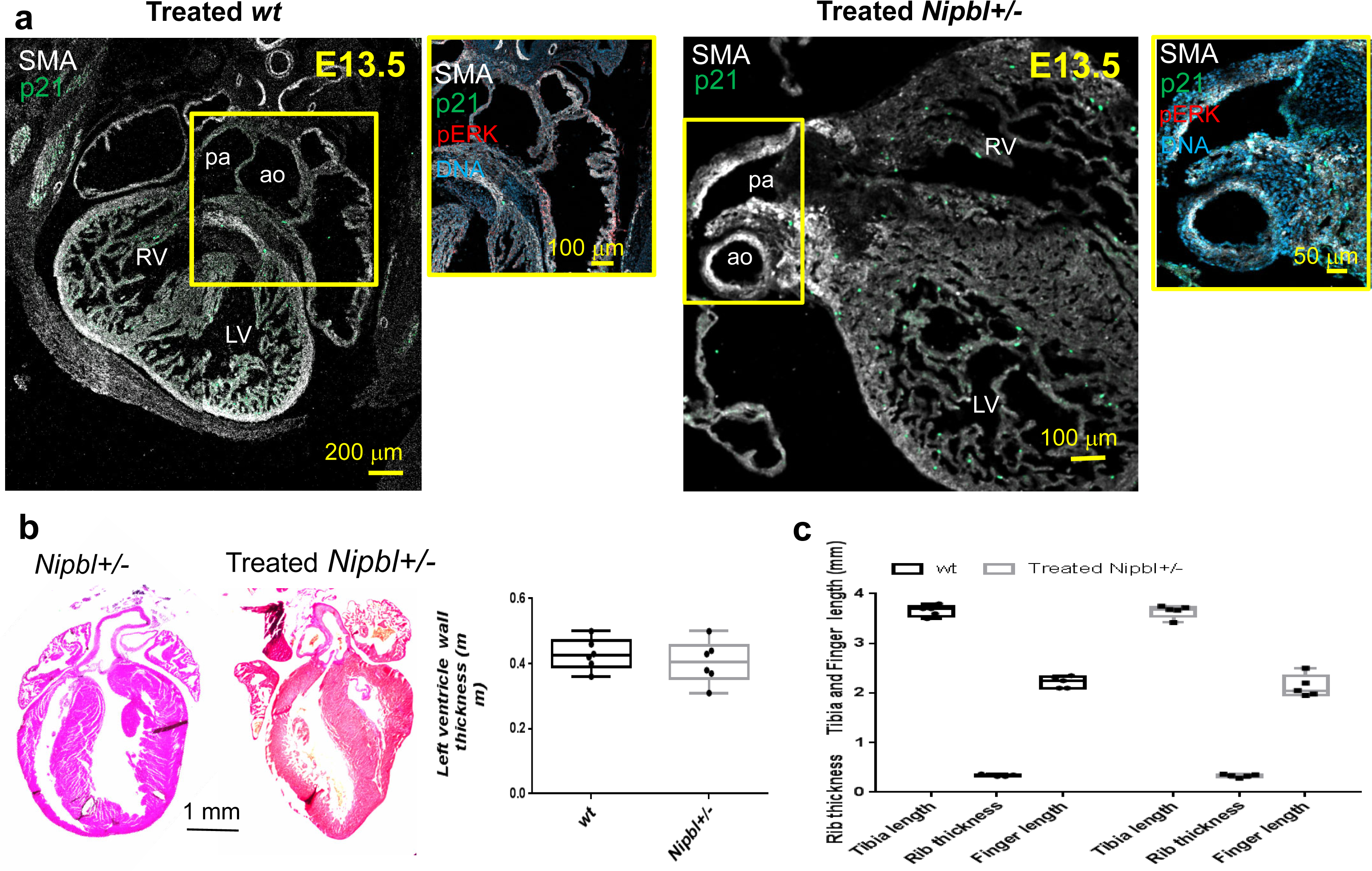
**Treatment of pregnant mice with Galunisertib, an ALK5 inhibitor rescues cardiac phenotype of *Nipbl+/-* offspring**. Pregnant mice were treated with 30 mg/kg galunisertib from E9.5 to E13.5 and embryos and hearts collected at E13.5 and stained with anti-SMA, -P21 and –PERK antibodies. (**a**), or left up to the delivery (**b,c**) (n=6).Neonatal hearts were stained with eosin-hematoxylin and left ventricular wall thickness was scored in 6 wt and 6 treated *nipbl+/-* hearts (**b**). **c**, bone lengths were measured in *wt* and *nipbl+/-* neonatal mice born from galunisertib-treated mothers.

Furthermore, the thickness (32± 5μm, n=3) of second heart field derived skeletal oesophagal muscle was fully restored in adult offspring from a galunisertib-treated mother and no or very few p21+ cells were detected in the muscle (Fig.S8a). Similarly, the number of p21+cells that were scored in the cortex of neonatal *Nipbl+/-* mice born from galunisertib-treated others was not significantly different from the one scored in wt neonatal mice (Fig. S8 b,c) The aorta *of nipbl+/-* adult mice originally born from galunisertib-treated mothers did not feature stenosis anymore (Fig S9)

Finally, in order to assess the relevance of these findings in human cells, we treated CdLS patient-specific iPS cells-derived smooth muscle cells with 10 μM galunisertib during their differentiation together with TGFβ1 and PDGFβ for 6 days (Fig.S10). In contrast to non-treated cells, galunisertib treated smooth muscle cells did not express p21 (Fig S10), indicating that in humans as in mouse model, attenuation of TGFβ signaling pathway protects cells from senescence.

Altogether, these results demonstrate that pharmacological inhibition of TGFβ signaling pathway, which is found hyperactive in our CdLS mouse model, in pregnant mice fully restores cardiac function and body size in newborn mice and also prevents senescence in newborn mice and in CdLS iPSCs derived smooth muscle cells.

## Discussion

*Nipbl+/-* haplo-insufficient mice in the C57Bl/6J genetic background recapitulate many features of the CdLS patients. More specifically, cardiac hypertrophy, PTA, ventricular septal defect and aortic stenosis were observed in a large majority of mice. The mice were shorter and featured deficient skeletal muscle derived from the second heart field including the oesophagal muscle, the masseters, all muscles involved in food intake. This muscle phenotype could explain the gastro-oesophagal reflux observed in CdLS children, at least partly.

The ventricular hypertrophy was likely a consequence of the persistent truncus arteriosus affecting the hemodynamics in the outflow trunk following both a thickening of the vessel wall and stenosis^36^.

Several observations point to senescence of several cell types including smooth and skeletal muscle cells in *nipbl+/-* mice which limited both their number and their motility. Indeed, E13.5 *nipbl+/-* embryos featured many senescent cells at the base of the outflow trunk (Fig 4). OFT explant experiments further highlighted a decrease in motility of these cells at E10.5, i.e., at the onset of EMT of endocardial cells in the outflow trunk (Fig S7) This observation likely accounts for a septation defect of the OFT.

The smooth muscle cells at the base of the outflow trunk derive from progenitors of the second heart field^37^ and possibly from a myocardial to smooth muscle cell trans- differentiation^38^. This explains that deletion of *Nipbl* in neural crest cell lineage did not affect heart and more specifically OFT formation^29^.

A good candidate to account for the senescence of smooth muscle cells is TGFβ2. Indeed TGFβ family has been involved in senescence of many cell types^39^. More specifically TGFβ2 is an inducer of smooth muscle cell senescence^40^, consistent with the protection of cells against senescence conferred by TGFβ inhibition we report in this work.

Smooth muscle cells at the base of the outflow trunk deficient in *Nipbl+/-* mice are regulated by both TGFβ2 and retinoic acid pathways, two pathways dysregulated in *Nipbl+/-* mice. Overactivation of TGFβ2 associated with a decrease in RA signaling^36^, a frequent association of events^41,42^, lead to defect in septation of OFT^37^.

TGFβ2 availability in OFT is likely increased in *Nipbl+/-* mice by a decrease in expression of extracellular matrix proteins including fibrillin, known to sequester the growth factor^33^.

The OFT phenotype could have been further worsened by a collagenolytic activity of senescent smooth muscle cells, a phenomenon mediated by the Senescence Associated Secretory Phenotype (SASP), and regulated by p38 MAPK^43^. More generally senescence is associated with a dysregulation of the extracellular matrix (ECM)^40^. This may account for the downregulation of many ECM proteins we observed in the proteomics profile of the E16.5 embryonic OFT of *Nipbl+/-* mice while they remained unaffected at the RNA level. The fact that many genes were upregulated while many proteins were downregulated in *Nipbl+/-* mice also point to a block in translation likely due to defect in nucleoli (Fig 6) and in turn in ribosomal activity.

We also observed senescence of both undifferentiated and differentiated smooth muscle cells from CdLS patients (Fig 6). Finally, cell senescence was also present in smooth muscle cells of adult *Nipbl+/-* aorta (Fig 3) as well as in the second heart field derived skeletal oesophagal muscle, in hypertrophic chondrocytes and in neonatal brains (Fig. S5).

The regulation of embryonic senescence of the growth plate cartilage by the TGFβ superfamily was reported as a determinant of the length of the bones^441^. An exacerbated TGFβ pathway induced cell senescence in primordium cartilage (Fig 4c) in *Nipbl+/-* mice likely explains their delayed growth. Along the same line, impairment in neuron migration was reported in *Nipbl+/-* neonatal brain^45^, which may also be due to a senescence of neurons just as the one we observed in neonatal brains.

Using a TGFβRI, ALK5 inhibitor, used in oncology (Galunisertib) in pregnant mice at mid-gestation in a time window between E9.5 and E13.5 rescued the size of the neonate (Fig.7, S7), restored the septation of the OFT in *Nipbl+/-* mice (Fig.7) and prevented cell senescence in skeletal oesophagal muscle, aorta, and brain cortex (Fig.S8,S9). The prevention of cell senescence in brain cortex might be clinically relevant as post-natal brain cortex features changes in chromatin 3D structure^46^ and still performs synaptogenesis^47^. This opens a postnatal therapeutic window.

The treatment of mothers also significantly decreased the number of senescent cells surrounding or in the OFT (Fig.7). *NIPBL+/-* IPS cell-derived smooth muscle cells treated with Galunisertib also prevented their senescence (Fig.S10). This further suggests that TGFβ2 mediates its deleterious function through cell senescence of many cell types.

Thus, collectively our data revealed that an exacerbated TFGβ pathway and associated cell senescence are at the origin of many defects in a CdLS mouse model. As this pathway is druggable, our research opens the path toward potential preventive, or even curative (at least for neuronal defects) therapeutic strategies for postnatal CdLS patients,

## Methods

### Mice

SMA*^creERT2+/-^* mice were obtained from Dr Daniel Metzger (IGBMC, Strasbourg). *Nipbl^flox/flox^* mice were obtained from Dr Heiko Peters, Institute of Genetic medicine, Newcastle University, UK. Mice were kept under standardized conditions (20–25°C temperature; 50%±20% humidity) on a 12 h light/12 h dark cycle in an enriched environment (kraft paper enrichment). Food and tap water were provided ad libitum. *Nipbl+/-* mice were generated by deleting exon 2 of the gene using *Nipbl^flox/flox^* mouse, bred with a CAG*^CreERT2+/-^* (JAX laboratory) mouse and gavaged with tamoxifen. The mice were then backcrossed for 10 generations in C57Bl6j genetic background. *Nipbl+/-* mice were maintained as heterozygous mice by breeding them with C57Bl/6J mice. Pregnant mice were separated from the male as soon as a vaginal plug was observed and daily observed. Two to three days before the female gives birth, a wild type C57Bl/6J mouse was added to the cage to prevent the mother from killing the small *Nipbl+/-* neonate. *Nipbl+/-* neonates were weaned after 4 weeks.

Tamoxifen (20 μg/g mouse) was given to SMA*^creERT2+/-^/Nipbl^flox/+^*female mice bred with *Nipbl^flox/flox^* (2 to 4 months old) males by oral gavage at E11.5.

Mice were genotyped by PCR of tail biopsies lysed in proteinase K for 3 hours using the primers of Table 1.

**Table 1:**
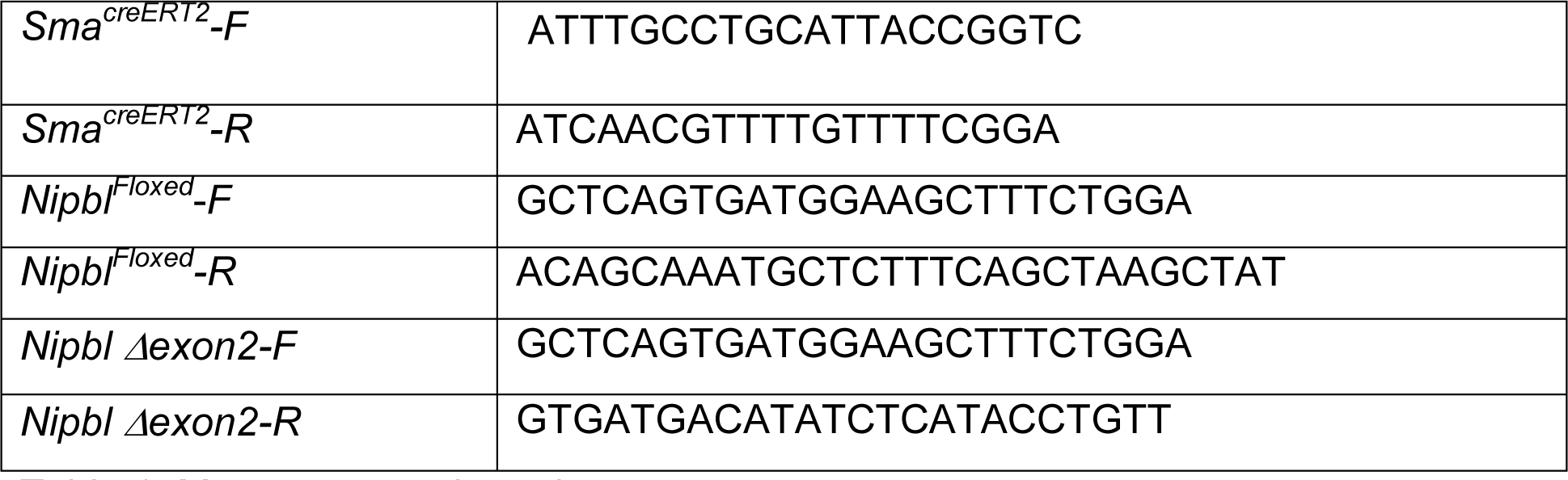
Mouse genotyping primers.

### High Resolution Episcopic Microscopy

Neonatal hearts of wt or *Nipbl+/-* were fixed for 24h in 4% PFA and washed in PBS before being embedded in plastic resin for HREM according to the manufacturer.

Images were processed using Imaris software.

### M-Mode and Doppler Ultrasound Transthoracic Echocardiography

Echocardiography was performed using an Affiniti 50 (Philips) and a 15 MHz probe. Mice were anaesthesized with 4% isoflurane and maintained with 1-1.5% isoflurane. Contractility of the left ventricle was acquired in a long-axis configuration in M-Mode. Aortic flow was monitored using PW Doppler-mode, by positioning the Doppler sample volume parallel to flow direction, assisted by Color Doppler-mode. Echocardiography recordings were analysed blinded by a cardiologist who had not performed the recordings. A minimum of 6 mice in groups were monitored.

### Antibodies

Antibodies used for cell or embryo immunofluorescence were raised against αSMA (Abcam, ab7817, 1/100), CD31 (Pecam, anti-mouse Pharmingen #550274 1/100), sarcomeric actinin (Sigma-Aldrich France ascites fluid EA-53, used at 1/2000), or p21 (Abcam Ab107099, 1/300), anti-pERK (santa-cruz sc-377400) anti-NIPBL (Bethyl A301-779A) used at 1/200 in immunofluorescence and in Immunoprecipitation (1 μg) and anti-N-terminal domain of NIPBL (given by JM Peters IMP, Vienna) used in western blot (1/1000) . Anti-SMC1was obtained from Abcam (ab9262).

### Cell imaging

Images were acquired using a confocal LSM 800 Zeiss microscope equipped with an airryscan of 32 detectors. Light was provided by a laser module 405/488,561 and 640 nm wavelengths, respectively.

All images were acquired using the ZEN ZEISS software. Then some images were deconvoluted using Autoquant and reconstructed in 3D using Imaris software (IMARIS). Episcopic images were also reconstructed in 3D using Imaris software.

Some large images (whole embryonic section, brain sections) were acquired using an Axioscan Z1 (Zeiss).All samples were mounted in Fluoromount™ (Cliniscience, France).

### Alzarin Red and Alcian Blue staining of neonatal skeleton

Skin and viscera were removed from sacrificed neonatal mice. The skeletons were then place for 5 days in Ethanol 95% and 2 days in acetone at room temperature. They were the stained with Alizarin Red/Alcian Blue for 5 days at 37°C. After a quick rinse in distilled water, they were placed in 1% KOH for 4 days and then in 20% glycerol/1% KOH for 5 more days. The skeletons were imaged using an Axiozoom V.16 and a 0.5x/0.125 FWD 114mm PlanAPo Z objective (Zeiss).

### Eosin and hematoxylin and MOWAT staining of hearts

Cryosections of hearts were stained with eosin and hematoxylin according to standard procedures or with MOWAT pentachrome stain kit (Biosite MPS2) according to manufacturer instruction.

### In situ hybridization and Safranin Weigert staining of forelimbs

Forelimbs were fixed in 4% formalin overnight. E18.5 and P0 forelimbs were decalcified in 25% EDTA overnight prior to embedding in paraffin. 5 µm serial sections were stained with Safranin Weigert and used for in situ hybridization using Digoxigenin-11-UTP labelled antisense probes against *Col2*, *Col10*, *Ihh* and *p21* as described previously (Schroeder et al. 2019). Brightfield images were taken on a Zeiss Axioplan2 microscope with a SPOT 14.2 camera and measures using SPOT advanced software. Fluorescent images were taken on a Zeiss Axio Observer7 microscope with an AxioCam 506 mono CCD camera and Zen2.3 software. Expression domains of all genes were measured using ImageJ. For each measurement, 3 to 6 sections were analyzed per animal. Length differences were compared between littermates.

### iPS cells culture and differentiation

IPS cells were derived from skin fibroblasts using Cytotune Sendai virus kit (Thermofisher, France) and cultured on MEF in KO-DMEM supplemented with KO-SR (Thermofisher, France), 10^-7^M mercaptoethanol, Non-Essential Amino Acids and 10 ng/ml FGF2 as previously described^48^. Cells were screened for any chromosomal defects using the iCS-digital TM PSC 24-probes kit (Stemgenomics, Montpellier, France) Cells were transferred on geltrex-coated plate in StemFit4 medium (NipponGenetics, Europe, Germany) for two passages and were differentiated into smooth muscle cells in RPMI-B27 medium supplemented with CHIR9901 8 μM for 24hrs, CHIR9901 4 μM and BMP2 10 ng/ml for 24hrs, BMP2 10 ng/ml and IWR1 5 μM for 24 hrs and for 6 days with TGFβ1 10 ng and PDGFβ 20 ng/ml.

### Proximal Outflow tract explants

Outflow tracts were dissected out from E10.5 wt and *nipbl+/-* embryos. The proximal part was cut and open along the long axis to be transferred on a collagen gel^493^, the endocardial side being on the gel. The explants were cultured for 48 hrs before being fixed with PFA 4%, permeabilised with Triton X100 0.1% and stained with DAPI. The explants were imaged in confocal microscopy using a 10X long distance objective. A z-stack of images was acquired. Distance of cell migration was scored using Image-J.

### RNA extraction, real time PCR and RNA sequencing

Total RNA was extracted from E16.5 OFT hearts (*Nipbl+/-* n=12 and controls n=12 from 6 separate litters) using Zymo Research Corp kit ZR RNA Miniprep following the manufacturer’s protocol. For the real time PCR, one µg of total RNA was used to synthetize cDNAs using oligo(dT) primers and affinity script reverse transcriptase (Agilent technologies France). Real-time quantitative PCR analyses were performed using the Light Cycler LC 1.5 (Roche, France). For each condition, expression was quantified in duplicate, and GAPDH was used as the housekeeping gene or normalizing RNA in the comparative cycle threshold (CT) method^44^. The sequences of *TGFβ1 sense GCTAATGGTGGACCGCAACAACG*, and *anti-sense CTTGCTGTACTGTGTGTCCAGGC*, for *TGFβ2 senseCACCTCCCCTCCGAAAATGCCAT,* and *anti-sense ACCCCAGGTTCCTGTCTTTGTGGT for* mouse *Nipbl*.

For the RNA sequencing, total RNA was isolated from E16.5 OFT hearts (*Nipbl+/-* n=12 and controls n=12 from 6 separate litters) and pooled either for *wt* or *Nipbl+/-* groups. The two pooled samples were used for the RNA-seq library preparation, using the kit TruSeq Stranded mRNA by Illumina.

Libraries were paired-end sequenced on the Illumina NextSeq 500 sequencer. Reads with a phred score lower than 20 and shorter than 25 bp were removed using Sickle (v1,33). Quality of trim reads was checked using multiQC (v1.0). Trim reads were aligned using STAR aligner (v2.7.0d) with arguments “outFilterMismatchNoverLmax” and “outFilterMultimapNmax” set to 0.08 and 1, respectively.

Transcripts discovery was performed using Cufflinks (v2.2.1) with the “library-type” argument set to fr-firstrand, and a GTF file obtained from GENCODE (“Comprehensive gene annotation”, vM1) provided as the genomic annotation. The GTF files produced for each sample by Cufflinks were combined using Cuffmerge. The “class code” assigned to each transcript by Cuffmerge was used to defined unknown transcripts (class code“u”). Only de novo transcripts with counts greater than 0 in at least one RNA-seq sample were kept for subsequent analyses. These de novo transcripts were combined with the GENCODE GTF file to produce the final genomic annotation that was provided to Feature Counts (v1.6.1) for quantification.

Differential gene expression was performed using DESEQ2 between MVDD and CTRL. To create bigwig files, reads from Watson and Crick strands were selected using SAMtools (v1.9) and provided to the bam2wig.py script from the RseQC program suite (v2.6.4).

### Proteomics

OFT were dissected out from E16.5 hearts (*Nipbl+/-* n=4 WT n=4 from 2 separate litters) and processed as individual samples. Samples were first run in SDS-PAGE stained with silver to normalize protein content. 3 µl out of 15 µl were injected in triplicate into the Mass spectrometer Orbitrap Fusion Lumos on a EasySpray ES803a-rev2 column of 50cm. Data were analyzed using MAXQUANT software.

### Western blots and immunoprecipitations

Protein extracts were separated by 10% sodium dodecyl sulfate–polyacrylamide gel electrophoresis (SDS-PAGE) and transferred onto nitrocellulose membranes (Invitrogen). Blocking and antibody incubations were performed in 5% bovine serum albumin. Membranes were incubated with HRP-conjugated anti-mouse or anti-rabbit secondary antibodies (Jackson ImmunoResearch) 1h at room temperature. Antibody detection reactions were developed by enhanced chemiluminescence (Millipore).

### Statistics

Results are presented as +/- SEM or as median and min to max The Student’s t test was used to analyze statistical significance. All p values corresponded to a two-tailed test, and a p <0.05 was considered statistically significant (**p<0.01; *p<0.05). The Graphpad software (Prism) was used to perform statistical tests.

## Acknowledgments.

This study was funded by the E-RARE European program COHEART.

We are grateful to the Leducq Foundation for generously awarding us for cell imaging facility (MP “Equipement de Recherche et Plateformes Technologiques” (ERPT).

We Thank Fabrice Prin (The Francis Crick Institute, London) for processing the neonatal hearts for HREM and for image acquisition and Chris Hunter (Indigo, London) for help in management of these HREM experiments, Drs Luc Camoin and Stéphane Audebert (Centre de Recherche en Cancérologie de Marseille, CRCM Inserm UMR1068, CNRS UMR7258, Aix Marseille Université U105) for expert processing and analysis of the samples in proteomics, Pr Juan Pié (Unit of Clinical Genetics and Functional Genomics, School of Medicine, Universidad de Zaragoza, Spain) for providing cells from CdLS patients and Dr Thomas Moore-Morris (INSERM U1251, Marseilles) for generating the *Nipbl+/-* mouse.

## References

1. Kline, A.D. et al. Diagnosis and management of Cornelia de Lange syndrome: first international consensus statement. Nat Rev Genet 19, 649–666 (2018).

2. Cascella, M. & Muzio, M.R. Cornelia de Lange Syndrome. in *StatPearls* (Treasure Island (FL), 2021).

3. Chatfield, K.C. et al. Congenital heart disease in Cornelia de Lange syndrome: phenotype and genotype analysis. Am J Med Genet A **158A**, 2499–505 (2012).

4. Selicorni, A. et al. Clinical score of 62 Italian patients with Cornelia de Lange syndrome and correlations with the presence and type of NIPBL mutation. Clin Genet 72, 98–108 (2007).

5. Buckingham, M., Meilhac, S. & Zaffran, S. Building the mammalian heart from two sources of myocardial cells. Nat Rev Genet 6, 826–35 (2005).

6. Krantz, I.D. et al. Cornelia de Lange syndrome is caused by mutations in NIPBL, the human homolog of Drosophila melanogaster Nipped-B. Nat Genet 36, 631–5 (2004).

7. Gao, D., Zhu, B., Cao, X., Zhang, M. & Wang, X. Roles of NIPBL in maintenance of genome stability. Semin Cell Dev Biol 90, 181–186 (2019).

8. Zuin, J. et al. A cohesin-independent role for NIPBL at promoters provides insights in CdLS. PLoS Genet 10, e1004153 (2014).

9. Sarogni, P., Pallotta, M.M. & Musio, A. Cornelia de Lange syndrome: from molecular diagnosis to therapeutic approach. J Med Genet 57, 289–295 (2020).

10. de Lange, C. Sur un type nouveau de degenciration (typus Amstelodamensis). . Arch Med Enfants 36:713-719. 36, 713-719. (1933).

11. Mills, J.A. et al. NIPBL(+/-) haploinsufficiency reveals a constellation of transcriptome disruptions in the pluripotent and cardiac states. Sci Rep 8, 1056 (2018).

12. Garcia, P. et al. Disruption of NIPBL/Scc2 in Cornelia de Lange Syndrome provokes cohesin genome-wide redistribution with an impact in the transcriptome. Nat Commun 12, 4551 (2021).

13. Pistocchi, A., et al. Cornelia de Lange Syndrome: NIPBL haploinsufficiency downregulates canonical Wnt pathway in zebrafish embryos and patients fibroblasts. Cell Death Dis 4, e866 (2013).

14. Santos, R. et al. Conditional Creation and Rescue of Nipbl-Deficiency in Mice Reveals Multiple Determinants of Risk for Congenital Heart Defects. PLoS Biol 14, e2000197 (2016).

15. Kawauchi, S. et al. Multiple organ system defects and transcriptional dysregulation in the Nipbl(+/-) mouse, a model of Cornelia de Lange Syndrome. PLoS Genet 5, e1000650 (2009).

16. Muto, A., Calof, A.L., Lander, A.D. & Schilling, T.F. Multifactorial origins of heart and gut defects in nipbl-deficient zebrafish, a model of Cornelia de Lange Syndrome. PLoS Biol 9, e1001181 (2011).

17. Kawauchi, S. et al. Using mouse and zebrafish models to understand the etiology of developmental defects in Cornelia de Lange Syndrome. Am J Med Genet C Semin Med Genet 172, 138–45 (2016).

18. Gu, W. et al. Defects of cohesin loader lead to bone dysplasia associated with transcriptional disturbance. J Cell Physiol (2021).

19. Singh, V.P., McKinney, S. & Gerton, J.L. Persistent DNA Damage and Senescence in the Placenta Impacts Developmental Outcomes of Embryos. Dev Cell 54, 333–347 e7 (2020).

20. Olley, G. et al. Cornelia de Lange syndrome-associated mutations cause a DNA damage signalling and repair defect. Nat Commun 12, 3127 (2021).

21. Vrouwe, M.G. et al. Increased DNA damage sensitivity of Cornelia de Lange syndrome cells: evidence for impaired recombinational repair. Hum Mol Genet 16, 1478–87 (2007).

22. Ball, A.R., Jr., Chen, Y.Y. & Yokomori, K. Mechanisms of cohesin-mediated gene regulation and lessons learned from cohesinopathies. Biochim Biophys Acta 1839, 191–202 (2014).

23. Lindsay, M.E. & Dietz, H.C. Lessons on the pathogenesis of aneurysm from heritable conditions. Nature 473, 308–16 (2011).

24. Chetaille, P. et al. Mutations in SGOL1 cause a novel cohesinopathy affecting heart and gut rhythm. Nat Genet 46, 1245–9 (2014).

25. Little, P.J., Tannock, L., Olin, K.L., Chait, A. & Wight, T.N. Proteoglycans synthesized by arterial smooth muscle cells in the presence of transforming growth factor-beta1 exhibit increased binding to LDLs. Arterioscler Thromb Vasc Biol 22, 55–60 (2002).

26. Gil-Rodriguez, M.C. et al. De novo heterozygous mutations in SMC3 cause a range of Cornelia de Lange syndrome-overlapping phenotypes. Hum Mutat 36, 454–62 (2015).

27. Lima, B.L. et al. A new mouse model for marfan syndrome presents phenotypic variability associated with the genetic background and overall levels of Fbn1 expression. PLoS One 5, e14136 (2010).

28. Vignier, N., Mougenot, N., Bonne, G. & Muchir, A. Effect of genetic background on the cardiac phenotype in a mouse model of Emery-Dreifuss muscular dystrophy. Biochem Biophys Rep 19, 100664 (2019).

29. Smith, T.G. et al. Neural crest cell-specific inactivation of Nipbl or Mau2 during mouse development results in a late onset of craniofacial defects. Genesis 52, 687–94 (2014).

30. Diogo, R. et al. A new heart for a new head in vertebrate cardiopharyngeal evolution. Nature 520, 466–73 (2015).

31. Comai, G. et al. A distinct cardiopharyngeal mesoderm genetic hierarchy establishes antero- posterior patterning of esophagus striated muscle. Elife 8(2019).

32. Heude, E. et al. Unique morphogenetic signatures define mammalian neck muscles and associated connective tissues. Elife 7(2018).

33. Godwin, A.R.F. et al. The role of fibrillin and microfibril binding proteins in elastin and elastic fibre assembly. Matrix Biol 84, 17–30 (2019).

34. Makki, N. & Capecchi, M.R. Cardiovascular defects in a mouse model of HOXA1 syndrome. Hum Mol Genet 21, 26–31 (2012).

35. Laforest, B., Bertrand, N. & Zaffran, S. Genetic lineage tracing analysis of anterior Hox expressing cells. Methods Mol Biol 1196, 37–48 (2014).

36. Buffinton, C.M., Benjamin, A.K., Firment, A.N. & Moon, A.M. Myocardial wall stiffening in a mouse model of persistent truncus arteriosus. PLoS One 12, e0184678 (2017).

37. Li, P., Pashmforoush, M. & Sucov, H.M. Retinoic acid regulates differentiation of the secondary heart field and TGFbeta-mediated outflow tract septation. Dev Cell 18, 480–5 (2010).

38. Liu, X. et al. Single-Cell RNA-Seq of the Developing Cardiac Outflow Tract Reveals Convergent Development of the Vascular Smooth Muscle Cells. Cell Rep 28, 1346–1361 e4 (2019).

39. Zhang, Y., Alexander, P.B. & Wang, X.F. TGF-beta Family Signaling in the Control of Cell Proliferation and Survival. Cold Spring Harb Perspect Biol 9(2017).

40. You, W. et al. TGF-beta mediates aortic smooth muscle cell senescence in Marfan syndrome. Aging (Albany NY) 11, 3574–3584 (2019).

41. Chen, F. et al. Inhibition of Tgf beta signaling by endogenous retinoic acid is essential for primary lung bud induction. Development 134, 2969–79 (2007).

42. Kubalak, S.W., Hutson, D.R., Scott, K.K. & Shannon, R.A. Elevated transforming growth factor beta2 enhances apoptosis and contributes to abnormal outflow tract and aortic sac development in retinoic X receptor alpha knockout embryos. Development 129, 733–46 (2002).

43. Balint, B. et al. Seno-destructive smooth muscle cells in the ascending aorta of patients with bicuspid aortic valve disease. EBioMedicine 43, 54–66 (2019).

44. Lui, J.C. et al. Differential aging of growth plate cartilage underlies differences in bone length and thus helps determine skeletal proportions. PLoS Biol 16, e2005263 (2018).

45. van den Berg, D.L.C., et al. Nipbl Interacts with Zfp609 and the Integrator Complex to Regulate Cortical Neuron Migration. Neuron 93, 348–361 (2017).

46. Tan, L. et al. Changes in genome architecture and transcriptional dynamics progress independently of sensory experience during post-natal brain development. Cell 184, 741–758 e17 (2021).

47. Zeiss, C.J. Comparative Milestones in Rodent and Human Postnatal Central Nervous System Development. Toxicol Pathol 49, 1368–1373 (2021).

48. Neri, T. et al. Human pre-valvular endocardial cells derived from pluripotent stem cells recapitulate cardiac pathophysiological valvulogenesis. Nat Commun 10, 1929 (2019).

49. Runyan, R.B. & Markwald, R.R. Invasion of mesenchyme into three-dimensional collagen gels: a regional and temporal analysis of interaction in embryonic heart tissue. Dev Biol 95, 108–14 (1983).

