## supplemental figures for "TGFβ−induced embryonic cell senescence at the origin of the Cornelia de Lange syndrome"

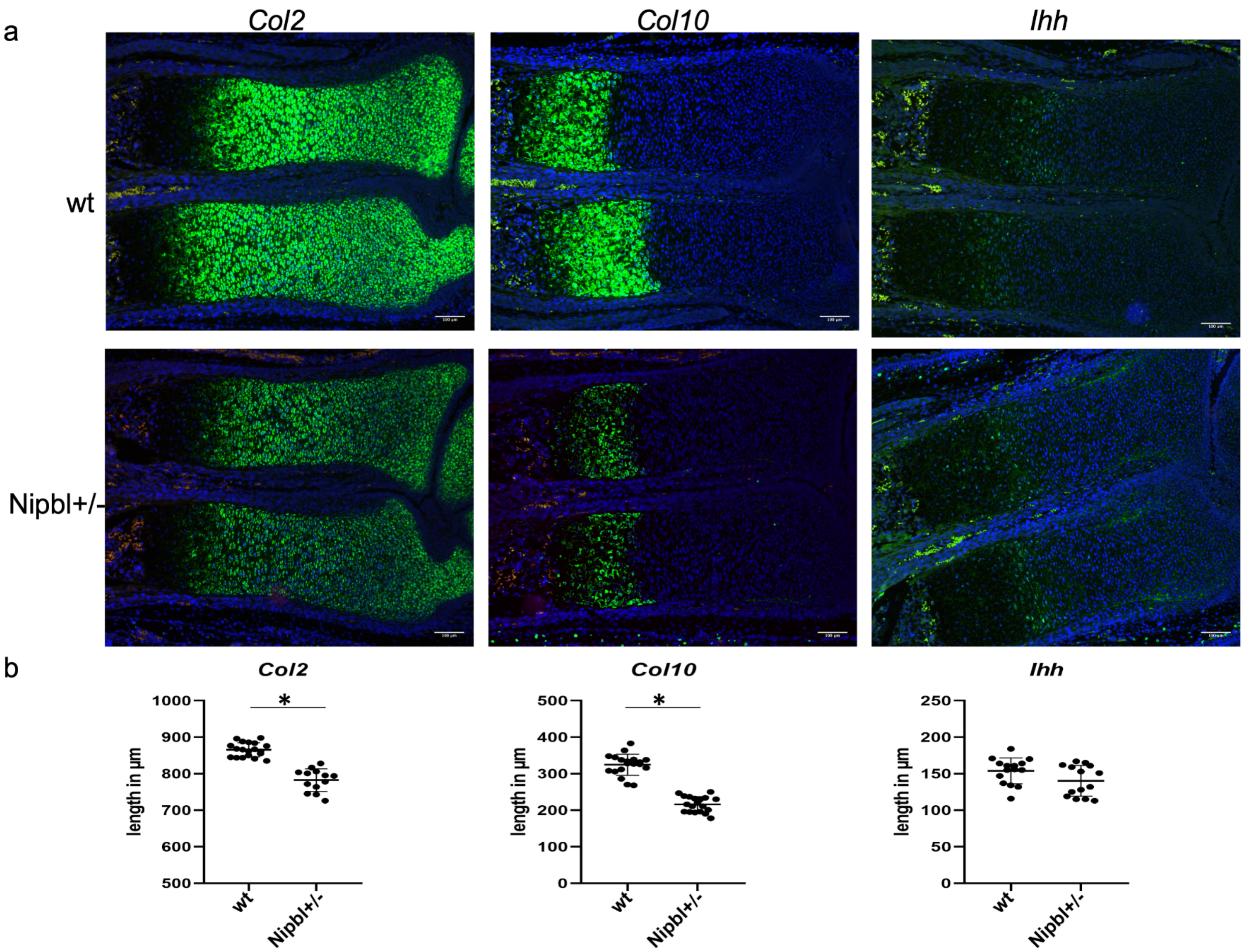


**Fig. S1 Reduced expression domains of Col2 in proliferating and *Col10* in hypertrophic chondrocytes of E16.5 *Nipbl+/-* mice. a**, The expression domains *of Col2*, *Col10* and *Ihh* were analysed by in situ hybridization in forelimbs section of *Nipbl+/-* and wt mice. Scale bar: 100 µm. **b**, The length (µm) of each expression domain was measured and plotted. Dots show individual measurements in several bone sections from 2 *Nipbl+/-* and 2 *wt* embryos *p<0,05).

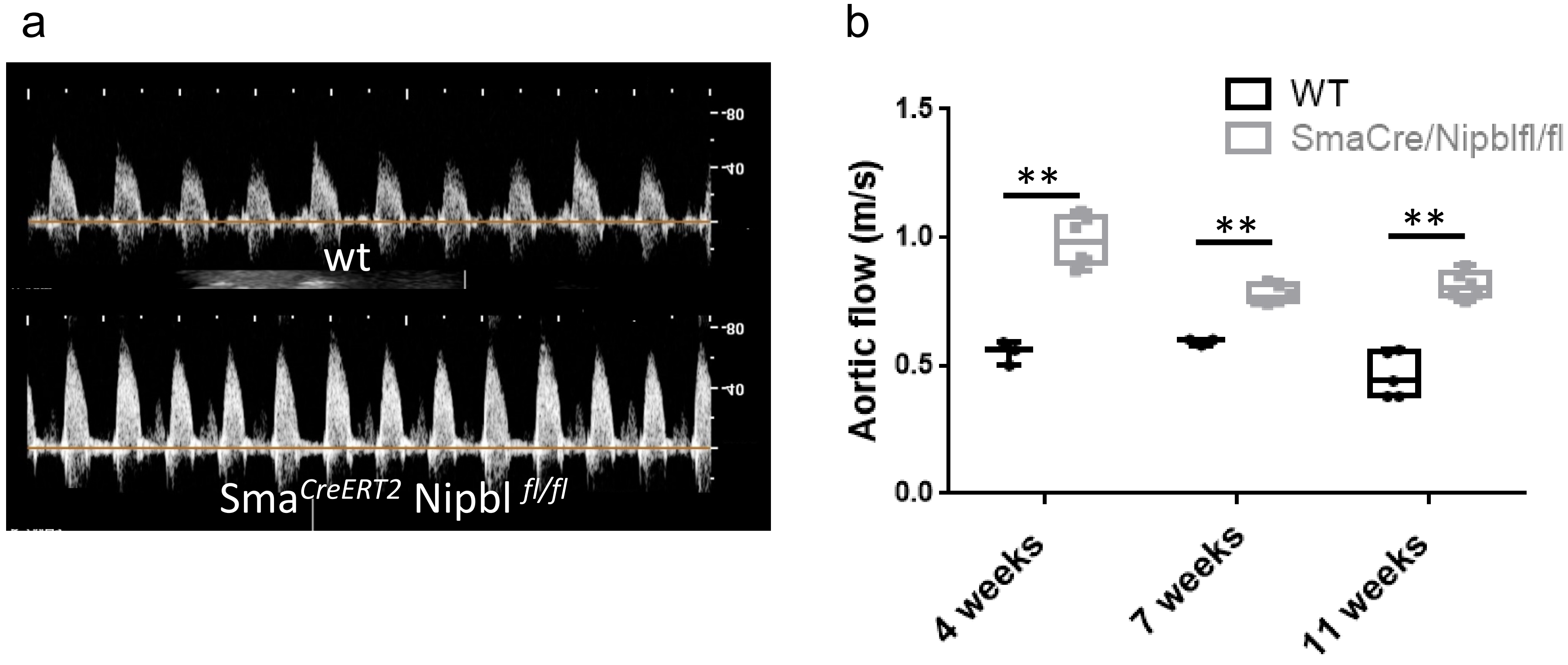


**Fig. S2 Conditional deletion of *Nipbl* specifically in the smooth muscle cell lineage** using the SMA$^{CreERT2}$ mouse recapitulates **a)** increases in aortic flux as monitored in Doppler echocardiography **b,** aortic flows scored at 4, 7 and 11 weeks p= 5 mice **t-test p<0.001

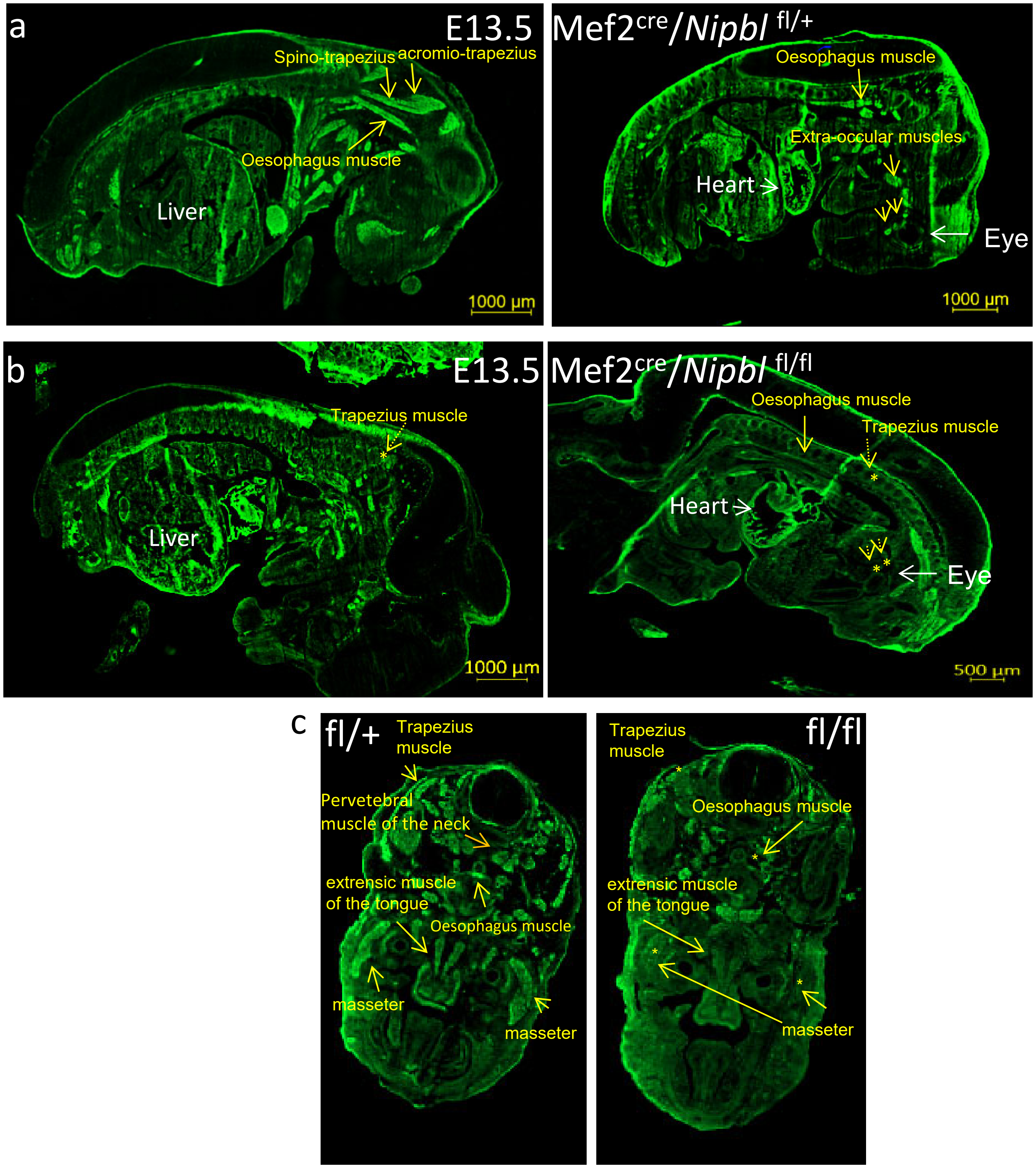


**Fig. S3 Defects in development of second heart field derived skeletal muscles in *Nipbl* haploinsufficient mice**
**a**,**b** Anti-actinin immunostaining of embryonic muscle saggital (**a**,**b**) and **c**, transversal sections of E13.5 Mef2c$^{Cre}$/*Nipbl* $^{fl/+}$ (**a**) and Mef2c$^{Cre}$/Nipbl $^{fl/fl}$ (**b**) mouse *embryos*
The yellow arrows point for the specific 2nd heart field derived skeletal muscles

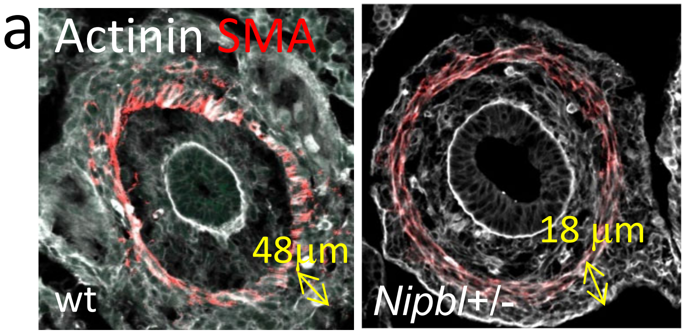


**Fig. S4 Cell senescence in skeletal oesphagal muscle, bone and brain cortex a,b,** actinin, SMA and p21 immunostainings of sections of oesophagus in 2 months old mice, **c**, of cortex of neonatal brains from wild type (wt) and *Nipbl+/-* mice. Insets show high magnifications of representative cortex area stained with an anti-p21 antibody. The images are respresentative of 4 wt and 4 *Nipbl+/-* mice.

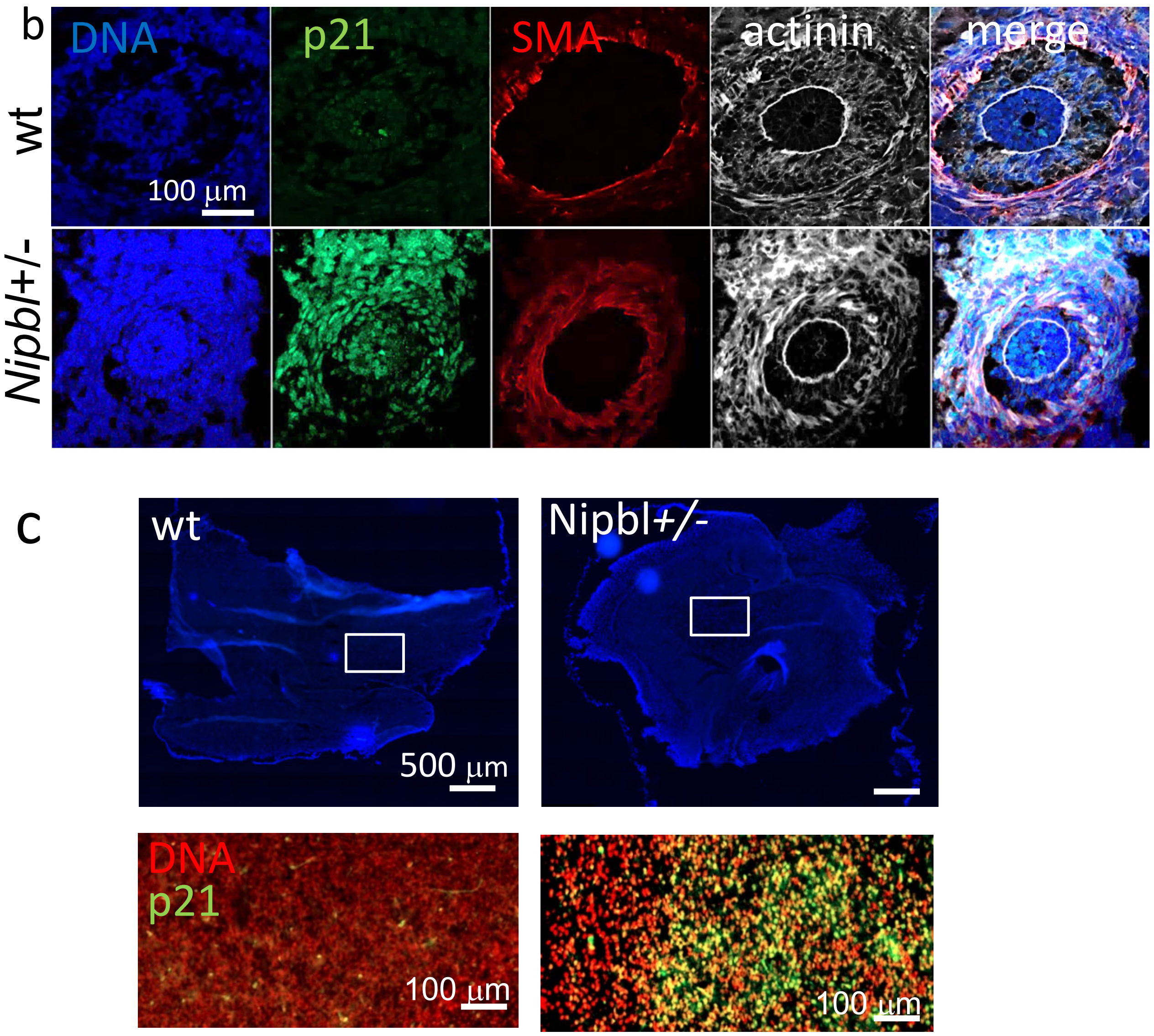

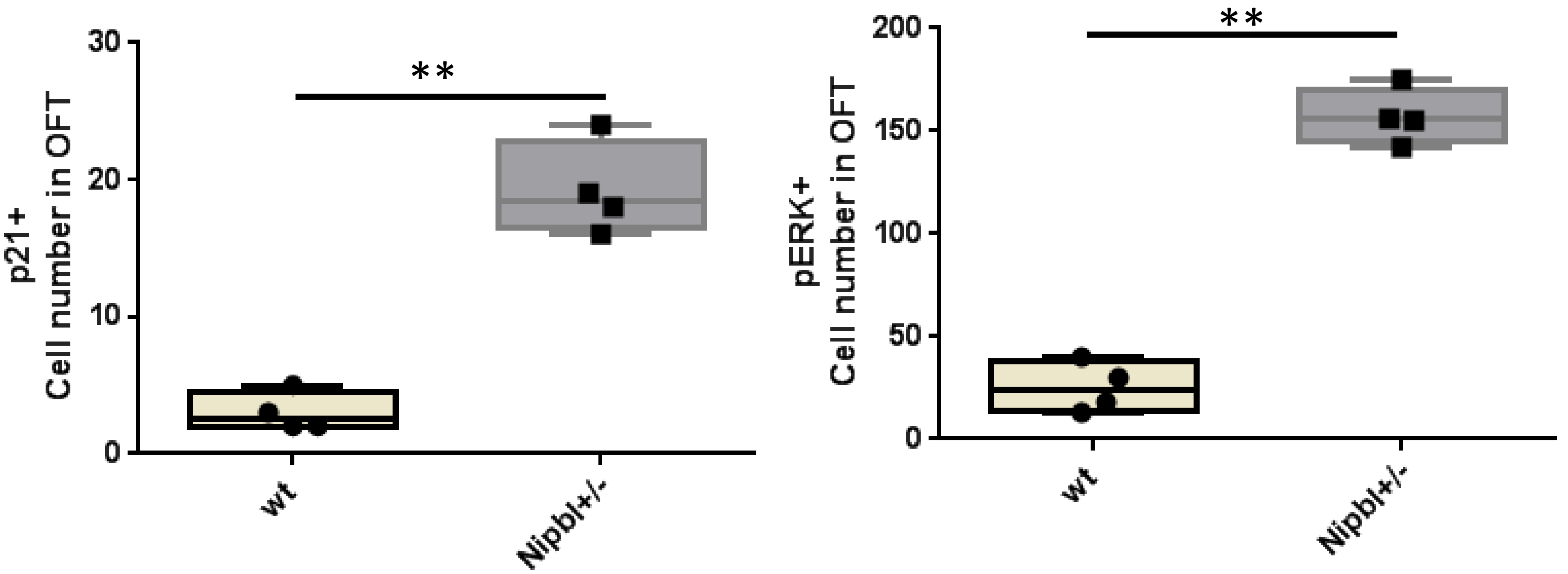


**Fig. S5 Numbers of p21+ (a) and pERK+ cells in OFT of wt and *Nipbl* +/- E13.5 OFT.**
Cells were scored by Imaris (nuclear spots) in a surface of 0.002 $cm^2$ using dapi staining as a reference. **t-test $p<0.001$

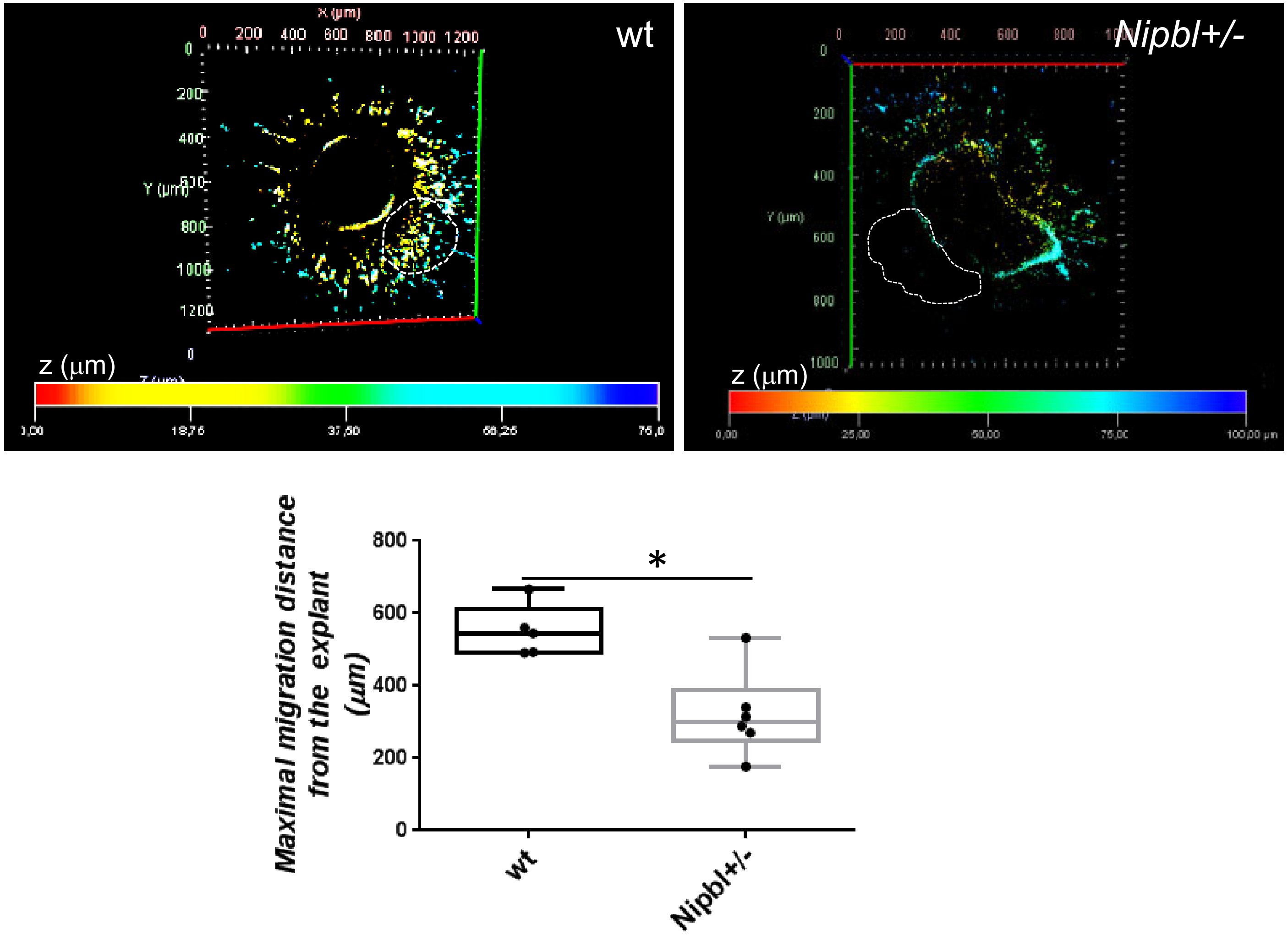


**Fig. S6 Invasive properties of proximal OFT cells.** OFT of the proximal ventricular region were dissected out from E10.5 embryonic hearts from *wt* or *Nipbl+/-* mice and cultured on collagen gels. After 48hrs cell migration from the explants was scored in 3D. Wt cells migrated farther and deeper into the gel than *Nipbl+/-* cells . The images are representative of 6 experiments ( 6 OFT explants/condition from 4 litters) *t-test $p<0.01$

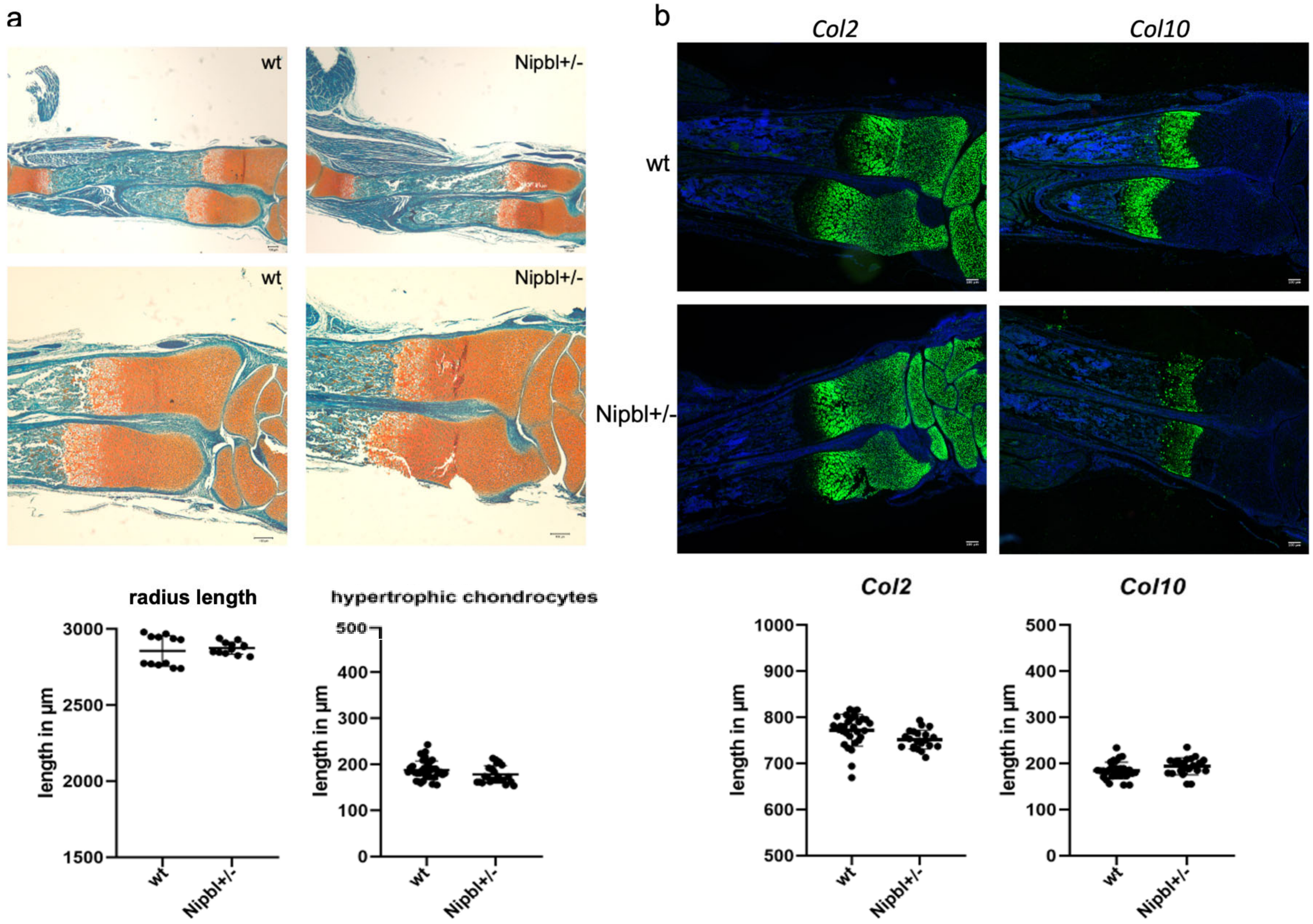


**Fig. S7 Galunisertib treatment rescued the skeletal phenotype of *Nipbl+/-* mice**.
**a**,Total radius length and the hypertrophic zone were measured in Safranin-Weigert stained forelimb section of galunisertib treated neonate N*ipbl+/-* and wt mice. Both zones showed similar length in both genotypes. **b**, In situ hybridization of *Col2* and *Col10 revealed similar expression domains i*n galunisertib treated neonate *Nipbl+/-* and wt forelimb sections. Dots show individual measurements of serial sections from 2 *Nipbl+/-* and 3 *wt mice*.

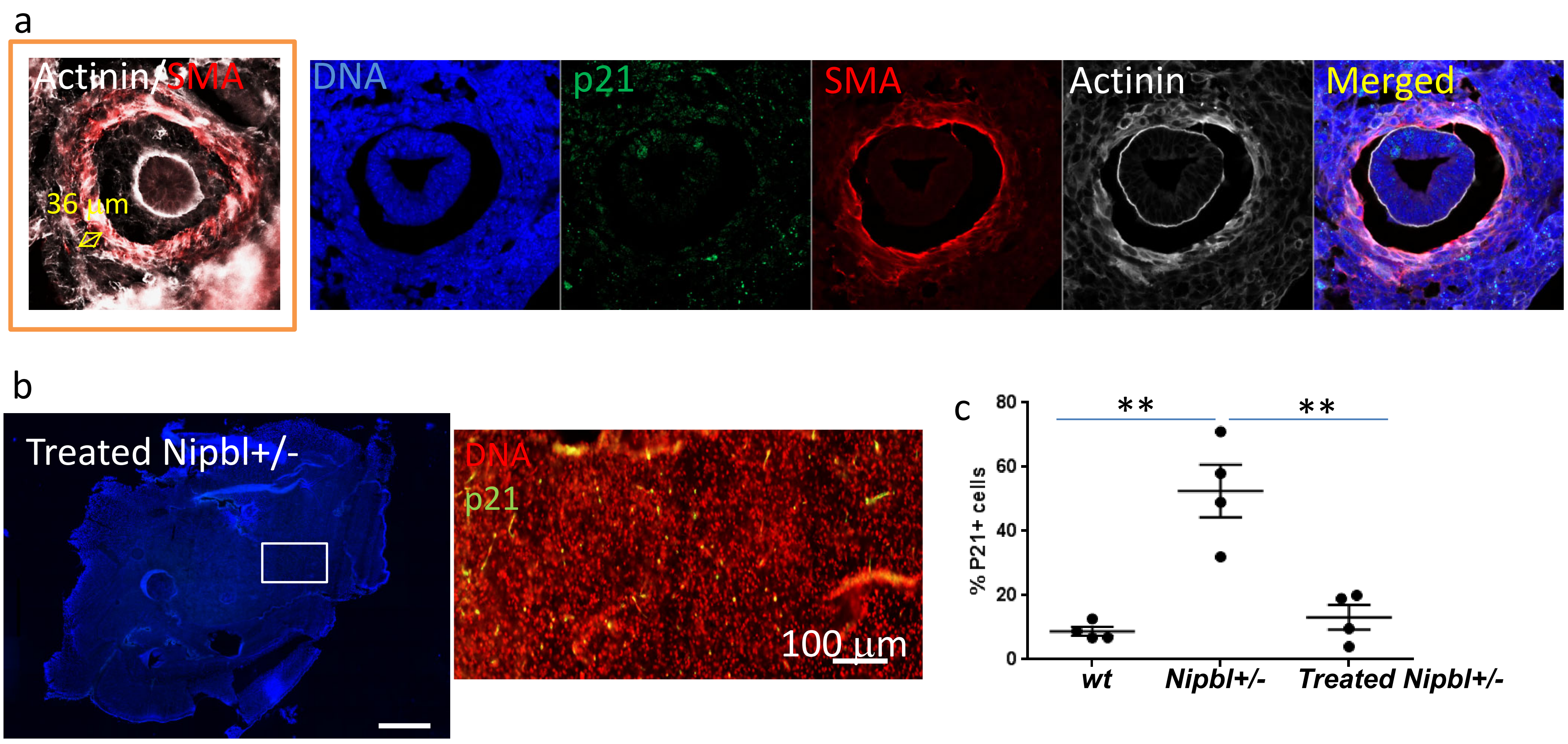


**Fig S8: galunisertib prevents cell senescence in both skeletal muscles and brains. a,** SMA, p21, actinin immunostaining of sections of oesophagus in 2 months old mice and **b**, of cortex of neonatal brains from *nipbl+/-* mice born from mothers treated with galunisertib diring gestation. Inset (a) measurement of skeletal muscle layer of the oesophagal muscle.**c**, scoring of p21+ cells in *wt*, *Nipbl+/-* and treated *Nipbl+/-* neonatal brains , normalized to Dapi+ nuclei ( n=4 mice, * t-test $p<0.001$).

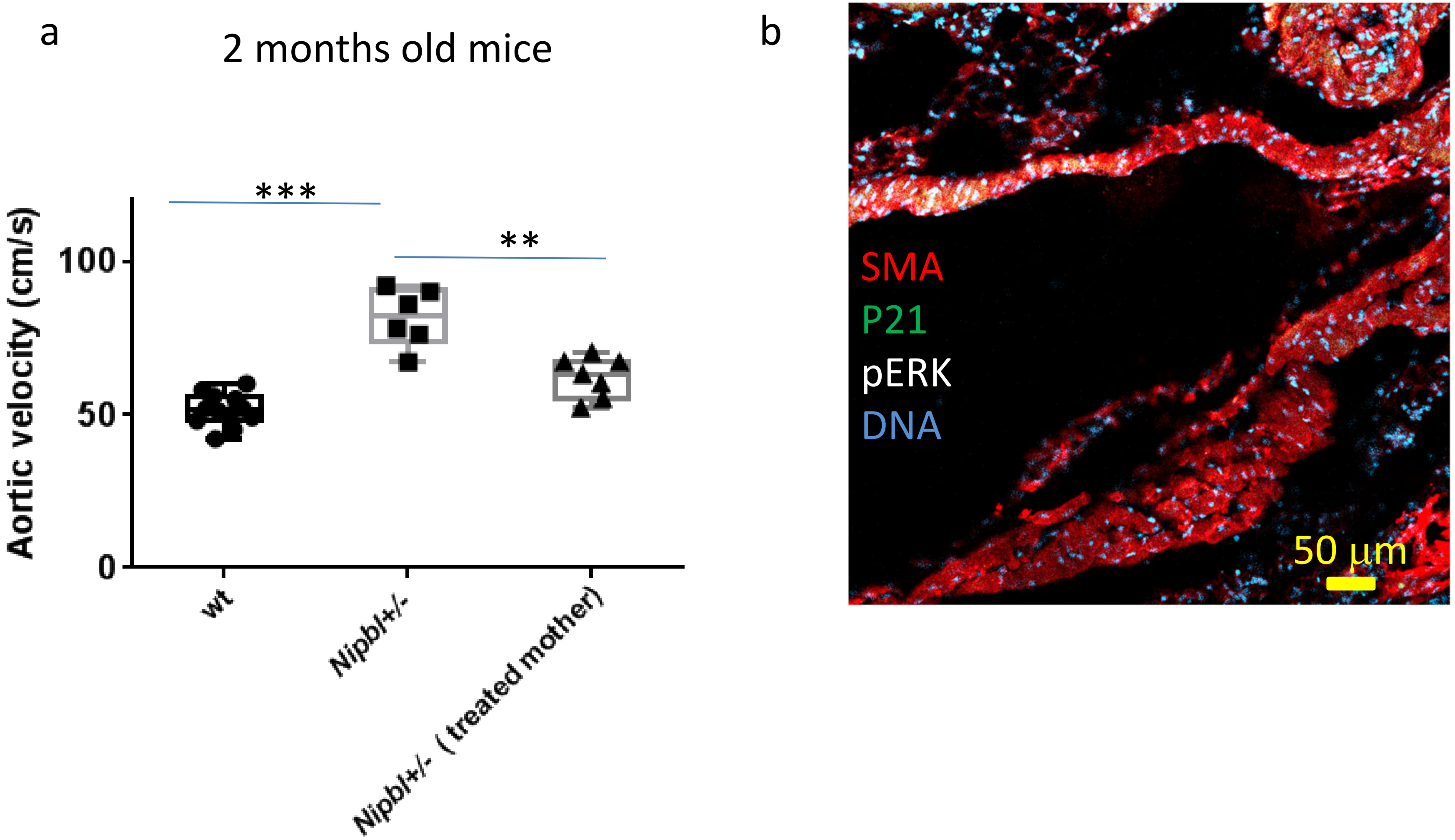


**Fig. S9 treatment of mother with Galunisertib prevents development of aortic stenosis** monitored by **a**, doppler echocardiography and **b**, aortic cell senescence (p21 and pERK staining of adult aorta) in offspring having reached adulthood. n=7 t-test 0.003***<p<0.017** ,

CdL iPS cell derived smooth muscle cells

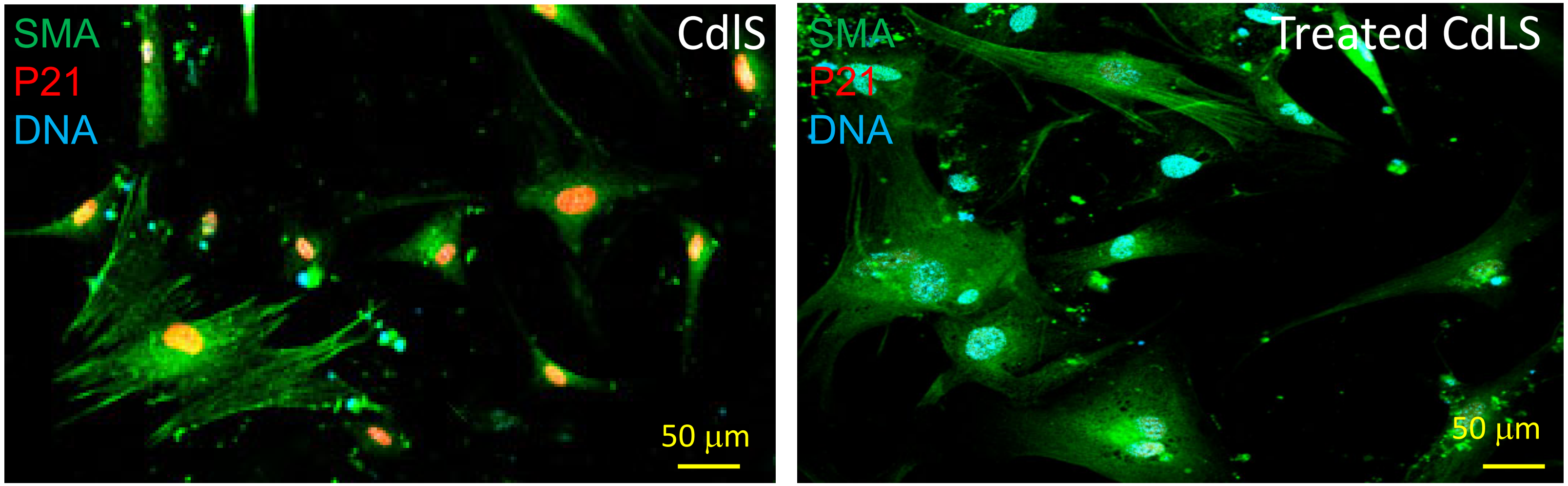


**Fig. S10 Treatment of smooth muscle cells from a CdlS patient with Galunisertib, an ALK5 inhibitor prevents senescence of NIPBL haploinsufficient cells.**
